# Naturalistic narrative listening reveals a core brain network for detecting moral information

**DOI:** 10.64898/2026.09.04.749401

**Authors:** Michal Klincewicz, Silvy HP Collin

## Abstract

Moral cognition is a complex process, which recruits a wide range of mechanisms. It is not clear whether any of them reliably reflects moral content in the stimulus. We set out to test whether a specific mechanism of moral content representation and/or detection exists, and where it is, if it exists. To that end we used uni- and multi-variate analyses, including inter-subject pattern correlation, on functional MRI recordings of people listening to naturalistic narratives with moral content. We discovered a core brain network for detecting moral content, which prominently involves the TPJ, MTG/STG, among other regions. Moral content is reliably detected across narratives and participants, independent of instructions to do so, prominently involving the TPJ that functions as a detector. We identify specific regions for moral features, such as vices/virtues. Our methods and results open a path to further understand how the brain processes specific features of moral content.

## 1. Introduction

Moral decision-making is a complex process known to involve a number of brain regions and networks both during the active decision-making role and also during the passive processing of morally-laden stimuli (Sevinc and Spreng, 2014). Most studies that aim to provide a mechanistic account of moral decision-making involve an isolated decision with several options presented. For example, participants are asked to make a choice between passively letting five hypothetical people get run over by a runaway trolley or to actively divert the trolley to run over one person (Crockett, 2013; Amit and Greene, 2012). From this literature, we know that especially lateral frontal regions, such as the dorsolateral prefrontal cortex (dlPFC) and the ventrolateral prefrontal cortex (vlPFC), as well as other regions of the executive network, are primarily involved in evaluating morally salient decisions slowly and deliberately, weighing evidence and applying norms and principles (Cushman, 2013; Arrouet et al., 2026). This is sometimes described as System 2 thinking (Greene and Haidt, 2002). Medial and limbic regions, including the ventromedial prefrontal cortex (vmPFC) and amygdala, are associated with more intuitive, fast, and reactive moral decisions, such as those involving immediate harm (Cushman et al., 2012; Koenigs et al., 2007; Greene et al., 2001). Mentalizing, broadly supported by a network of regions including mPFC, hippocampus, temporoparietal junction (TPJ) and posterior cingulate cortex (PCC), also involve striatal regions in the context of decision making, which in turn facilitate imagery relevant to the decision at hand, including mentalizing, bringing episodes from one’s past up for comparison, and visualizing possible consequences of a decision (O’Connor and Fowler, 2023). Some studies focus on the processing of morally-laden information, independent of any decision. Here, a wide range of frontal (including the dorsomedial and ventromedial prefrontal cortex), parietal, and medial regions are implicated (Sevinc and Spreng, 2014).

One challenge for a unifying account of moral cognition that involves both the decision component and passive processing is that it is difficult to disentangle moral information from other aspects of the presented stimulus. Watching a movie, reading a vignette, or listening to a story all involve networks specific to processing that sort of content in general, independently of whatever moral content there may be contained therein. When moral information is operationalized by contemporary psychological theories of morality, such as Moral Foundations Theory (MFT) (Chen et al., 2026) or Morality as Cooperation Theory (MAC) (Weber et al., 2024), the network of regions identified is also connected specifically to that theory.

In this study, we search for the existence of a core moral network that operates independently of specific task demands, stimulus content or explicit moral reasoning. We aim to identify a set of brain regions that consistently tracks morally-laden information across diverse narrative contexts, present during passive listening in the absence of explicit moral judgments or instructions. Secondly, we predict that different moral foundations may recruit additional neural systems over and above this core set of brain regions. Therefore, beyond identifying a core moral network, we investigate whether distinct moral content differentially engage regions over and above the core moral network regions.

## 2. Results

### 2.1. Core moral network

To identify the core moral network in the brain, we ran univariate wholebrain analysis, separately for the *Tunnel* story and the *21st year* story, and furthermore separately for vices and for virtues. Then, we ran a minimum conjunction analysis (Nichols et al., 2005) to determine which brain regions were consistently active across both narratives and across vices and virtues (for more details see Methods) which revealed clusters in bilateral middle and superior temporal gyri (MTG, STG) and the TPJ, along with sensorimotor areas including the precentral and postcentral gyri (Figure 1 and Table 1). Additional activations were observed in the precuneus, cuneus, and intraand supracalcarine cortex as well as in the anterior cingulate cortex (ACC) and paracingulate cortex. Given the conservative filtering applied, these results strongly suggest that a specific network of regions is recruited for processing moral narrative content.

**Figure 1.**
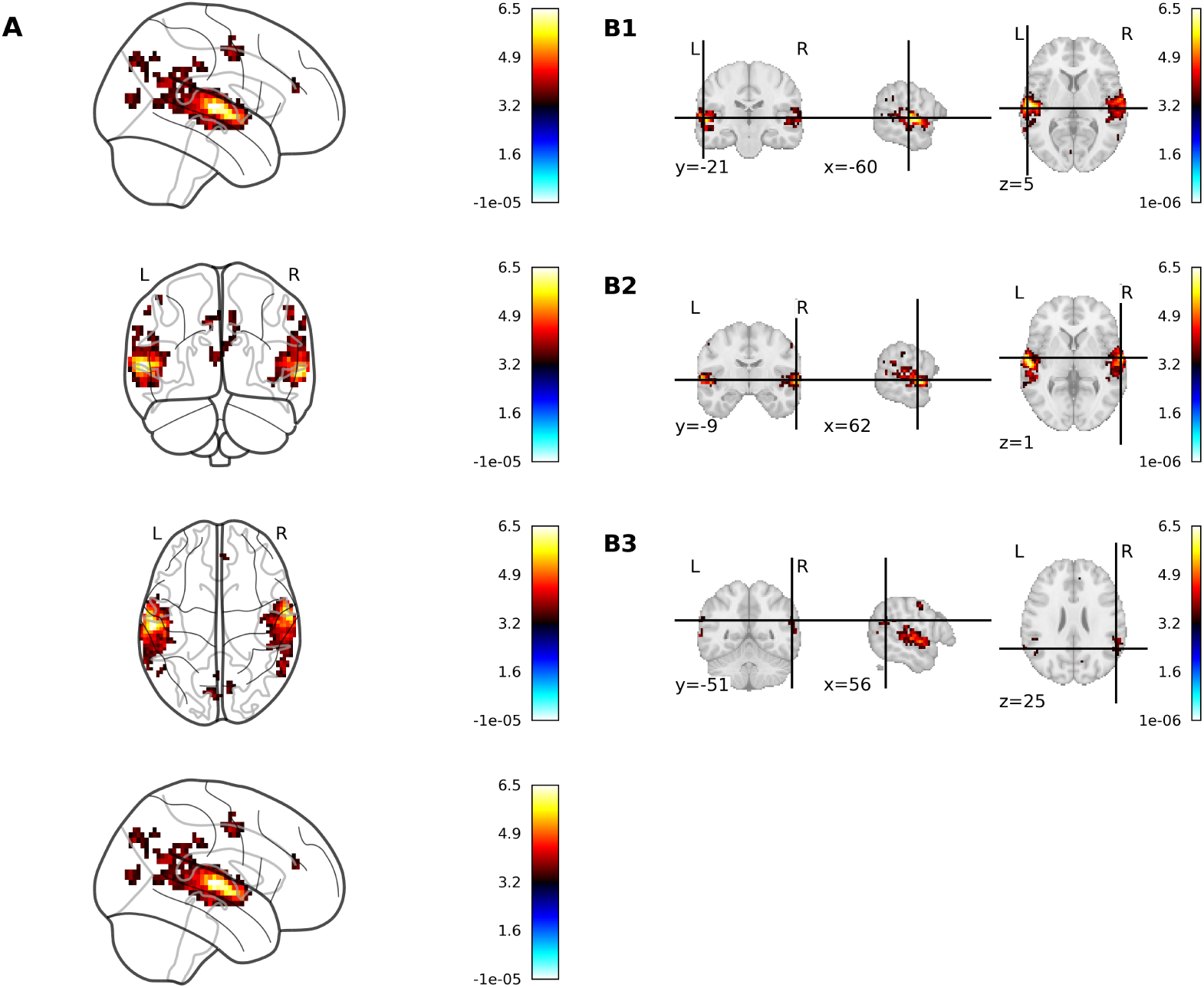
Core moral network. [A]. FDR-corrected clusters (*≥* 5 voxels) identified in the minimum conjunction analysis across the vice F-contrast and the virtue F-contrast, and across *Tunnel* and the *21st Year*. Major clusters are located in the MTG, STG, TPJ, pre-and postcentral gyrus, precuneus, cuneus, intra- and supracalcarine cortex, anterior cingulate cortex (ACC) and paracingulate cortex. **[B]** Slices for the three largest clusters, corresponding to left MTG/STG **[B1]**, right MTG/STG **[B2]** and TPJ **[B3]**. See Table 1 for more details.

**Table 1.** Core moral network regions. Cluster locations and anatomical composition for all suprathreshold clusters (*≥* 5 voxels) identified in the minimum conjunction analysis across the vice F-contrast and the virtue F-contrast, and across *Tunnel* and *21st Year*. Coordinates (*x, y, z*) are reported in MNI space. Mean F-values and cluster volumes (mm^3^) are shown for each cluster. Percentages reflect the proportion of each cluster falling within Harvard–Oxford atlas regions. Abbreviations: STG = Superior Temporal Gyrus; MTG = Middle Temporal Gyrus; COC = Central Opercular Cortex; SMG = Supramarginal Gyrus; AG = Angular Gyrus; PaCG = Paracingulate Gyrus; CG = Cingulate Gyrus; Cun = Cuneal Cortex; POC = Parietal Operculum Cortex; PreCG = Precentral Gyrus; PostCG = Postcentral Gyrus; HG = Heschl’s Gyrus; PCu = Precuneus; ICC = Intracalcarine Cortex; SCC = Supracalcarine Cortex; PT = Planum Temporale; PP = Planum Polare; p = posterior; a = anterior; s = superior, i = inferior; to = temporo-occipital.

| $x$ | $y$ | $z$ | mean<br>F | $\text{mm}^3$ | Harvard–Oxford atlas regions |
| --- | --- | --- | --- | --- | --- |
| -60.5 | -21.5 | 5.5 | 3.99 | 16272 | 23.67% Left p-STG; 17.04% Left PT;<br>10.40% Left COC; 10.40% Left a-STG;<br>9.96% Left p-MTG; 9.29% Left HG |
| 62.5 | -9.5 | 1.5 | 3.89 | 13968 | 30.93% Right p-STG; 17.27% Right<br>PT; 10.57% Right a-STG; 10.31% Right<br>COC; 8.51% Right HG; 6.70% Right<br>PP; 5.41% Right p-SMG |
| 56.5 | -51.5 | 25.5 | 3.62 | 1656 | 56.52% Right AG; 32.61% Right p-SMG; 6.52% Right sLOC |
| 56.5 | -6.5 | 45.5 | 3.56 | 684 | 89.47% Right PreCG; 10.53% Right<br>PostCG |
| -3.5 | -81.5 | 9.5 | 3.51 | 684 | 68.42% Left ICC; 26.32% Left SCC;<br>5.26% Right SCC |
| 8.5 | -75.5 | 41.5 | 3.44 | 576 | 81.25% Right PCu; 12.50% Right s-LOC; 6.25% Right Cun |
| -63.5 | -42.5 | 29.5 | 3.52 | 540 | 80.00% Left p-SMG; 20.00% Left a-SMG |
| 65.5 | -30.5 | 37.5 | 3.49 | 324 | 100.00% Right a-SMG |
| 2.5 | 44.5 | 17.5 | 3.44 | 288 | 87.50% Right PaCG; 12.50% Right a-CG |
| -6.5 | -78.5 | 37.5 | 3.59 | 288 | 62.50% Left PCu; 37.50% Left Cun |
| -54.5 | -3.5 | 49.5 | 3.54 | 288 | 87.50% Left PreCG; 12.50% Left<br>PostCG |
| 53.5 | -60.5 | 9.5 | 3.44 | 252 | 85.71% Right i-LOC; 14.29% Right to-MTG |
| -51.5 | -39.5 | 21.5 | 3.62 | 252 | 85.71% Left POC; 14.29% Left p-SMG |
| -51.5 | -57.5 | 25.5 | 3.46 | 252 | 57.14% Left s-LOC; 42.86% Left AG |

### 2.2. ISPC-based RSA reveals moral similarity structure in right TPJ

For the top 3 neural clusters (left MTG/STG, right MTG/STG, and TPJ) of the core moral network (see Figure 1B), we examined whether patterns of neural activity systematically reflected moral foundation scores using an inter-subject pattern correlation (ISPC) analysis (see Methods for more details). To do this, we first extracted left and right TPJ activation patterns and then created sentence-wise representational RDMs and compared these to the moral-foundation-score-RDM based on sentence-wise moral foundation score differences. The neural RDMs were computed using a leave-oneparticipant-out procedure (See Inter-subject pattern correlation (ISPC)).

RDMs from the TPJ were positively correlated with the moral-foundation-score-RDM for both stories (Left TPJ for *Tunnel*: Spearman *ρ* = 0.021; right TPJ for *Tunnel*: *ρ* = 0.025; left TPJ for *21st Year*: *ρ* = - 0.0005; right TPJ for *21st Year*: *ρ* = 0.0063). Although small in magnitude, permutation testing (i.e., 1000 random shuffles of the moral-foundation-score-RDM) confirmed that correlations for both stories were significant (right TPJ for *Tunnel*: p = 0.005; right TPJ for *21st Year*: p = 0.016; left TPJ for the *Tunnel* p = 0.053) and not significant in the left TPJ for *21st Year* (p = 0.554). See Appendix C for more details on the statistical analysis.

These findings suggest that the multi-voxel activation patterns across sentences in the right TPJ mirror the similarity structure of per-sentence moral foundation scores. In other words, sentences that were more similar to each other (i.e., similar moral-nonmoral ratio scores) evoked similar neural patterns, while morally different sentences produced more dissimilar neural activity patterns.

Although the magnitude of the correlations was modest (around 0.02)—as is typical in representational similarity analyses of high-dimensional fMRI data (Kriegeskorte et al., 2008)—the robust permutation-based significance indicates high reliability. We conclude that the right TPJ carries structured, content-sensitive information about moral meaning during passive listening to a narrative. The TPJ likely plays a representational role in encoding moral similarities or differences (See Discussion).

We did a similar ISPC analysis for the MTG/STG cluster. Here, the neural RDMs from the MTG/STG were positively correlated with the moral foundation score RDM for the *Tunnel* (left MTG/STG for *Tunnel*: Spearman *ρ* = 0.041, p-value based on permutation testing: p = 0.02; right MTG/STG for *Tunnel*: *ρ* = 0.062, p-value based on permutation testing: p = 0.006). In contrast to the (right) TPJ results, this did not replicate for left or right MTG/STG in the *21st Year* (left MTG/STG for *21st Year*: Spearman *ρ* = 0.007, p-value based on permutation testing: p = 0.08; right MTG/STG for *21st Year*: *ρ* = 0.006, p-value based on permutation testing: p = 0.163, Appendix C).

### 2.3. Differential neural engagement across moral domains

Section 2.1 describes the core moral network. Next, we were interested in assessing whether our fMRI data showed differential neural engagement for distinct moral foundations. To find out, we used T-contrasts that compare each moral foundation to the mean of the other 6 moral foundations (separately for virtues and for vices and separately for the *Tunnel* and *21st Year*). Table 2 (for *Tunnel*) and Table 3 (for *21st Year*) list which brain regions, over and above the core moral network regions, were significantly active for a given distinct moral foundation (FDR-corrected). Appendix D outlines the full table of FDR-corrected results for each of these T-contrasts.

**Table 2.** Brain regions that are not in the core moral network and that survive FDR-correction in a T-contrast comparing individual foundation *>* mean of other six foundations in the *Łuллel* story, separately for virtue and vice. Abbreviations: MFG = Middle Frontal Gyrus; SFG = Superior Frontal Gyrus; IFG = Inferior Frontal Gyrus; ITG = Inferior Temporal Gyrus; PCC = Posterior Cingulate Cortex; FPC = Frontopolar Cortex; PHG = Parahippocampal Gyrus; FO = Frontal Operculum; INS = Insula; TP = Temporal pole; AMY = amygdala; THAL = Thalamus; a = anterior

| Foundation | Virtue (additional) | Vice (additional) |
| --- | --- | --- |
| Deference |  |  |
| Fairness | INS; FO; FPC | AMY; PHG; THAL |
| Family | MFG; SFG; TP; IFG; **FPC**;<br>THAL; PHG; a-ITG | PCC |
| Group |  | FO; INS; AMY; IFG |
| Heroism |  |  |
| Property |  |  |
| Reciprocity |  |  |

**Table 3.** Brain regions that are not in the core moral network and that survive FDR-correction in a T-contrast comparing individual foundation *>* mean of other six foundations in the *21st Year* story, separately for virtue and vice. Abbreviations: INS = Insula; FO = Frontal operculum; FPC = Frontopolar Cortex; HPC = Hippocampus; PHG = Parahippocampal Gyrus; SPL = Superior parietal lobule; THAL = Thalamus; mPFC = medial prefrontal cortex; CAU = Caudate; LG = Lingual gyrus; PAL = Pallidum; FG = Fusiform Gyrus; PCC = Posterior Cingulate Cortex; OFC = Frontal orbital cortex; PUT = Putamen; NAcc = Nucleus Accumbens

| Foundation | Virtue (additional) | Vice (additional) |
| --- | --- | --- |
| Deference |  |  |
| Fairness |  | mPFC; FPC |
| Family | **<u>FPC</u>**; INS |  |
| Group | CAU; IFG; SFG; FPC | SPL |
| Heroism |  | FPC |
| Property | INS; FG; LG | SFG; MFG; FPC; INS; IFG;<br>THAL; CAU; PAL; HPC;<br>OFC; FG; PHG; PUT |
| Reciprocity | SFG; FPC; PCC; INS;<br>OFC; IFG; MFG; PUT;<br>NAcc; CAU; FG; LG; FO;<br>PAL |  |

Results suggest that fairness (virtue and vice), family (virtue and vice), and group (vice) foundations indeed have a distinct neural network compared to the other moral foundations in *Tunnel*. This largely replicated in *21st Year* as shown in Table 3. *21st Year* also showed a distinct neural pattern for fairness (in particular vice), family (in particular, virtue), and group (virtue and vice) compared to other moral foundations and a distinct network for property (virtue and vice) and reciprocity (virtue).

The foundation-specific activation patterns we uncovered do not closely resemble one another across the two narratives. The only point of commonality is the frontopolar cortex (FPC) in for the Family virtue foundation (bolded and underlined in Tables and 2 and 3. This is perhaps not surprising, given how dissimilar the narratives are in all respects (See Stimuli). First of all, as a story, we can stipulate that *21st Year* is not as engaging as *Tunnel*. It is, in fact, intentionally hard to follow and its auditory features are monotone due to a single narrator throughout. *Tunnel* is immersive and designed to keep a radio listener’s attention with distinct voice actors, lively dialogue, music, and sound effects. We see this reflected in our results, especially in Tables 2 and 3, where the activation of frontal regions as well as subcortical structures (e.g., thalamus, putamen) is much more prominent for the *21st Year* story. These regions are known to be important for vigilant attention (i.e., keeping sustained attention on the task) (Langner and Eickhoff, 2013). Besides that, frontal regions are also known to be prominently involved in directive attention and executive control (Rossi et al., 2009). In other words, participants listening to *Tunnel* in the scanner were having a much better time paying attention to the narrative and likely understanding it than those listening to *21st Year*. We provide further speculation on these differences in the Discussion, where we also offer a per-region interpretation in terms of individual foundations.

## 3. Discussion

In our study, we found a network of brain regions relevant for detecting moral content in general (i.e., core moral network) as well as distinct networks of brain regions responsible for specific moral foundations (e.g., fairness or group). Importantly, inter-subject pattern correlation analyses revealed that multi-voxel activation patterns in the core moral network (MTG/STG and TPJ) track the similarity structure of moral content within the story across participants. This strongly suggests that the core moral network supports a generalizable, stimulus-driven detection of moral content, not only in explicit moral judgments, but also during passive listening to morally-loaded narratives that can be substantially different from each other in content and form (See Stimuli). Additionally to this core moral network, we found networks of brain regions seemingly responsible for specific moral foundations, especially for fairness, family and group. We found these differ substantially depending on high-level features of the stimulus relevant to interpretability.

The core moral network (MTG and STG, TPJ, precentral and postcentral gyri, precuneus, cuneus, intra-and supracalcarine cortex, ACC and paracingulate cortex) has a detector-like function that involves specific downstream networks depending on the nature of the moral stimulus, including other high-level narrative features of the story. Our definition of the core moral network was very conservative: regions had to be consistently engaged across all moral foundations and for both virtue and vice. Contrary to what we expected and some of the current literature on moral cognition (Schuwerk et al., 2017; Koster-Hale and Saxe, 2013), the core moral network did not include frontal regions. Frontal regions were implicated in specific moral foundations (See below for details). One possible limitation of our conservative approach is that frontal regions may have been inadvertently excluded from the core moral network, especially if they are more selectively recruited.

All that said, according to our analysis, the most prominent regions within the core moral network are the MTG/STG and the TPJ (See Figure 1). MTG (Bzdok et al., 2012; Sevinc and Spreng, 2014; Chen et al., 2026) and STG (Khoudary et al., 2022; Chen et al., 2026) have previously been identified as a core region for moral decision making, empathy, and theory of mind. The MTG and STG are also primary sites for semantic processing and narrative comprehension (Binder and Desai, 2011; Babajani-Feremi, 2017). So it may be unsurprising that we find these regions in the core moral network. STG/MTG are likely processing moral meaning from the narrative.

The TPJ, on the other hand, has consistently been associated with moral cognition as part of the visual attention network and theory of mind network (Krall et al., 2015; Schuwerk et al., 2017; Koster-Hale and Saxe, 2013; Weber et al., 2024). A meta-analysis from 2012 (Bzdok et al., 2012) relates the TPJ (together with the MTG and dmPFC) as most consistently involved in empathy, theory of mind, and moral decision-making across all literature on these topics. While a lot of moral cognition literature is focused on active moral judgments, the TPJ has also been related to passive exposure to morally-laden stimuli (Sevinc and Spreng, 2014). The latter is more in line with the likely stipulated function of the TPJ in our study: as a detector. Besides knowledge on TPJ function in general, there is literature on lateralized functions of the TPJ which we will cover below when discussing left-right distinction in our multi-variate results.

Besides these prominent regions, the core moral network in our results also included both pre-and post-central gyri. The precentral gyrus (primary motor cortex) and the postcentral gyrus (primary somatosensory cortex) are primarily associated with motor and somatosensory processing, respectively, but are also prominently involved in embodied simulation (Gallese and Lakoff, 2005; Brecht, 2017). The presence of motor-related areas in moral reasoning has been described before, for example by Hopp et al. (2023) that describe the presence of the Supplementary Motor Area (SMA) in moral reasoning.

Also the precuneus is often mentioned in relation to moral reasoning (Schuwerk et al., 2017; Weber et al., 2024; Bzdok et al., 2012; Hopp et al., 2023) as well as social cognition (Khoudary et al., 2022; Alcalá-López et al., 2024). We interpret the presence of the precuneus in the core moral network to likely be related to mental imagery, episodic simulation and selfreferential processing, i.e., projecting the self into possible scenarios (Dadario and Sughrue, 2023). The strong presence of secondary visual regions (i.e., cuneus, intracalcarine cortex, supracalcarine cortex) strengthen the view that the participants are involved in visual imagery and mental scene construction of especially the morally relevant parts of the narrative.

The ACC has been linked to theory of mind (Schuwerk et al., 2017), empathy (Bzdok et al., 2012) and moral reasoning in general (Weber et al., 2024). Involvement of the ACC likely relates to the known role of the ACC in detection of valence, given it is part of the salience network (Alcalá-López et al., 2024; Arrouet et al., 2026). More specifically, the ACC has been linked to involvement in processing salient cues related to the self (Alcalá-López et al., 2024) and to emotional appraisal (O’Connor and Fowler, 2023). Its involvement in processing salient cues might be the reason why prior literature relates the ACC more to vice than to virtue (Chen et al., 2026).

Altogether, our results suggest a distributed core moral network that is reliably tracking moral information during passive listening (i.e., without explicit instructions to focus on moral content or without having to make explicit moral choices). This core moral network is centered around temporal regions (MTG, STG) and TPJ, but also includes midline and sensorimotor areas. This suggests that moral meaning is extracted spontaneously by a distributed network of brain regions when being exposed to morally-loaded content.

Following our mass univariate analysis approach to define which networks of brain regions were involved in processing the moral aspects of the narratives, we used a multi-variate pattern approach to shed more light onto the functioning of the most prominent regions within the core moral network. More specifically, we used inter-subject pattern correlation analysis (ISPC) to determine how the neural patterns within these core regions represent morally relevant vs morally irrelevant parts of the narrative across participants. Participants who process information similarly have more similar neural responses that can be captured by pattern recognition methods referred to as inter-subject correlation (as introduced by Hasson et al. (2004), but also see recent opinion article Ohad et al. (2025)). Statistical significance is then usually assessed using permutation testing (of e.g., 1000 iterations) to generate a null distribution against which the relevant correlation is tested. In our case, this approach was used to generate a null distribution reflecting no relationship between moral content and neural similarity (centered around zero, see Appendix C). Following prior work (Nguyen et al., 2019; Chen et al., 2017; Baldassano et al., 2017), p-values were calculated based on the mean and standard deviation of this distribution. Especially for subtle and context-sensitive shifts in sentence processing (as differences in moral load of sentences will likely be), ISPC is considered more sensitive than classical averaging-across-stimuli methods (Ben-Yakov et al., 2012). Using this method, we discovered that multi-voxel activation patterns in the right TPJ reflect the similarity structure derived from moral foundation scores based on the text of the stories (via MAC), for both stories (Tunnel and 21st Year). Thus, sentences in both of these stories with more similar moral profiles elicited more similar neural patterns in the right TPJ, preserved across participants. It has been shown before that the TPJ contains neural patterns for events preserved across participants (Baldassano et al., 2017; Nguyen et al., 2019). Our findings add to this that these shared event representations within the TPJ might, in part, represent moral meaning.

While the neural patterns within the right TPJ were shared more so for morally-loaded content across participants for both narratives (i.e., Tunnel and 21st year), the left TPJ approached significance in an ISPC analysis for the Tunnel story, but not for the 21st Year story. To speculate on this apparent difference across the narratives for the left TPJ, Krall et al. (2015) and Schuwerk et al. (2017) both relate the right TPJ in particular to basic cognitive functions like reorienting attention which is naturally relevant for processing moral content in any narrative. Looking at the MTG/STG, we found only a significant ISPC result for the Tunnel story, and not for the 21st Year story. We speculate that this distinction might be related to a difference in attention ability across the two narratives. The 21st year story might have led to neural patterns that are less well aligned across participants (that the ISPC analysis relies on) because the harder to follow narrative (i.e., 21st year) leads to more disengagement and more short shifts toward internally generated thought at different moments in time across participants compared to the Tunnel story. In other words, the ISPC results differed across the left TPJ and MTG/STG because of fluctuations in attentional demands generated by the stimulus. The engaging and shorter Tunnel story may have led most participants to attend externally, while the longer, more difficult 21st Year forced them inward. Given that the MTG and STG are both heavily involved in auditory and semantic processing (Petrides, 2023), it is not surprising to see in-between participant similarity across these regions suffer.

All 7 moral foundations rely on the core moral network. Are there additional regions necessary for processing moral content (virtue vs vice; specific moral foundations)? When looking at the results of the engaging Tunnel story, we see that virtue content engages frontal sub-regions, which perhaps reflecting abstract reasoning about principles (Morin et al., 2023), while vice additionally engages regions involved in salience detection or emotion, such as the amygdala and insula (parts of the salience network, Uddin (2016)), the PCC which has been linked to decision salience (Heilbronner et al., 2011) as well as processing emotional stimuli. These additional regions are popping out for fairness, family and group and not for the other moral foundations, but that might be because the Tunnel story most prominently relies on these foundations, not necessarily because in general fairness, family and group are different from the other foundations in terms of additional regions necessary. future work could manipulate which moral foundations a given story relies on to test this virtue vice distinction.

The 21st year results were different compared to the Tunnel story for these foundation specific effects, which we speculate is the case for the same reason as the distinction between the narratives as described for the ISPC results above. We speculate this has to do with the nature of the stimulus, that in case of the Tunnel story was designed to capture radio listeners attention and the 21st year was designed for research and needed to be two parallel stories in one overarching narrative that only later on gets properly integrated into one narrative. As a result, the constant switching makes it hard to follow which might have pushed people towards more shallow encoding of the meaning of the narrative (focusing on things like who belongs with whom, how to these locations and people all relate to each other). to actually engage with moral meaning of any given narrative, one might need to be more deeply immersed with the story then people did for 21st year. The most distinct difference that table 3 shows is the sudden recruitment of subcortical regions (striatum, caudate, putamen, pallidum,…), both on the vice and virtue side. This might have to do with the fact that these regions are known to be involved with sustained attention and focus on task relevant stimuli (Langner and Eickhoff, 2013), which is not directly related to moral content processing in that sense.

## 4. Conclusion

In this study, we show that some of the most abstract and controversial features of the social world–moral values–are objective in the sense that they can be perceived and cause selective activity in specific brain regions that function to represent them.

Our findings are consequential for neuroscience in at least two ways. First, we demonstrate that moral features in the environment are things we can be more or less good at detecting and understanding, depending on engagement. If we pick up moral cues in content, we represent moral content more fully. We can nevertheless detect that moral content is present in the narrative stimulus. This suggests that moral information processing is importantly subserved by a core moral network dedicated to both detection and representation. Second, we develop a method to identify networks of regions involved in processing specific features of moral content (virtue/vice, moral foundation, etc.) independently of decision-making.

## 5. Methods

### 5.1. Participants and fMRI dataset

All data were obtained from the open dataset “Narratives: fMRI data for evaluating models of naturalistic language comprehension” collection by Samuel A. Nastase and colleagues, deposited at the Princeton University open lab repository (https://datasets.datalad.org/labs/hasson/narratives/). The “Narratives” collection includes 28 naturalistic, unique stimuli that range from 3 to 56 minutes for a total of 5 hours of audio. 345 participants (204 female, 140 male, 18-53 years old, mean age 22) listened to one or more of these audio recordings while being in a scanner. Every narrative includes the audio file used during the recording and a text transcript. We used the fMRI data of 2 of these narratives, *Tunnel Under the World* (*Tunnel* from now on) and *21st Year*, with 48 participants (male/female, age) in sum. The main factor used to select the stimuli was the length of the recording, with a preference for longer audio.

### 5.2. Stimuli

*Tunnel* (N=23, 2 excluded; 25:34 minutes) is an old-fashioned radio audio narrative with voice actors. The story centers on a man who discovers that he is living in a simulated world. Format has lively dialogues, sound effects, and music. The narrative is immersive, using multiple voice actors for different characters, and designed to keep a radio listener’s attention. *21st Year* (N=25; 56:14 minutes) is two seemingly unrelated stories that eventually merge into one. Both revolve around an unhappy marriage. The audio is broken up by silences in between vignettes of interactions between members of the couples, a daughter, and a lover. It is narrated by a single narrator and harder to follow compared to the Tunnel story.

### 5.3. Identifying moral foundation values in the stimuli

We designed a multi-step natural language processing pipeline to identify the moral aspects of the two stories. First, each audio file was automatically transcribed with a speech-to-text model (Bain et al., 2023), which was then used to identify word onset and offset timestamps, which in turn identified sentence onsets and offsets in the audio with millisecond precision. Second, each sentence and chunk was scored using the extended MAC dictionary (Malik et al., 2025). We obtained a score for each foundation virtue and vice between 0 and 1 and a moral/non-moral score ranging from 0 to 7. The latter is an estimated strength of the moral signal in a sentence independent of individual foundation scores.

MAC identifies seven foundations separable by the nature of cooperation strategies in non-zero-sum games (See Table A.1). Each foundation corresponds to one such strategy (Curry et al., 2019):

### Family

Societies can coordinate for mutual benefit by valuing the allocation of benefits and harms to kin. So, in most societies, resources are altruistically allocated to one’s offspring and family at the expense of others, and doing so is deemed morally good. Not doing so or valuing non-kin over kin is, in many societies, deemed as morally wrong. In our stimuli, any narrative components related to kin likely scored higher in the family vice or virtue category.

### Group

Groups coordinate for mutual advantage on all levels of society and do so by creating institutions, hierarchies, and conventions, and then by enforcing boundaries for belonging. There are many ways in which this foundation manifests itself in our stimuli; whenever a social hierarchy or group is observed, discussed, or its effects are consequential.

### Reciprocity

Giving what you are owed in the context of social exchange of any kind, including employee-employer relations or revenge, are ways societies encourage and punish behaviors that affect group benefits. Reciprocity is found in norms governing proportionality of response, redistribution of wealth, but also punishment for littering, not paying for public transport, or cheating of any kind. Reciprocity may feature in narratives when there is an apology, gratitude, or forgiveness, but also a social moral emotion like guilt or shame.

### Heroism and Deference

Rank differences naturally generate conflicts. Societies developed ways of displaying dominance by proxies, such as athletic allegiance, cutting comments, or, in general, contests that signal bravery or ability–this is heroism. People can showcase their acceptance of their lower rank in relation to someone of a higher rank through rituals of humility and respect–this is deference. In narratives, these two foundations in social displays or all sorts, including witty comments or contests, such as battles, duels, or tense social gatherings.

### Fairness

Dividing resources is a happy compromise to open competition we may observe in the state of nature. Competition among members of a group is almost always a suboptimal way to get benefits that could have been gotten by distribution. Competition affects cohesion in a group and typically incurs costs to all parties involved, but especially those at the losing end of a competition. Fairness is a shorthand label for avoiding these costs.

### Property

Groups enforce norms around possession, which help avoid an open conflict. A possession or relationship is likely to be an instance of a property foundation in a narrative context.

### 5.4. MRI data acquisition and preprocessing

The fMRI dataset we used for this study used a 3T Siemens Magnetom Skyra scanner with a 20-channel head coil. Functional BOLD data were collected with interleaved gradient-echo EPI (TR/TE 1500/28 ms, 3×3×4 mm voxels, 27 axial slices, GRAPPA acceleration 2). We downloaded the already preprocessed data from the open source repository, which were volumetric fMRI-data in MNI152-space (see original article, Nastase et al. (2021), for details on the pre-processing steps).

### 5.5. Data analysis and statistics

#### 5.5.1. Univariate whole-brain fMRI analysis

##### First-level models

To identify the core moral network in the brain, we ran univariate whole-brain fMRI analysis. The functional MRI data were analyzed using two general linear models (GLMs) per story (i.e., 2 GLMs for the *Tunnel* and 2 GLMs for the *21st Year*), implemented in Python’s nilearn package (Abraham et al., 2014). We split modeling in that way because virtues and vices are highly confounded. The main point of difference, given how probability distributions for words and sentences are determined, is valence, which is computed in part using sentiment analysis (Malik et al., 2025).

A design matrix was constructed for the two GLMs using the task regressors of interest, modeled using the sentence-by-sentence moral foundation virtue score by using the onset timestamp and duration of the sentence. These durations were binned by foundation vice/virtue, giving 7 distinct task regressors of interest. The sentences that had zeros for all 7 virtue foundations were assigned to “baseline” (i.e., non-moral). The distribution of foundation scores across sentences is in Appendix B. For each of the two GLMs, additional nuisance regressors were: csf, white matter, translation x, y and z, and rotation x, y and z.

##### First-level contrasts

For each participant, we computed 7 T-contrasts for each of the two GLMs probing the effects of each individual regressor relative to baseline (e.g., virtue family regressor versus baseline), relative to the mean of the other 6 foundation scores (e.g., virtue family regressors versus the other 6 moral foundation regressors), and an additional F-contrast assessing the joint contribution of all 7 moral foundation regressors of the GLM.

##### Second-level model

Contrast images from the first level were used to create a second-level model for the group. For each T-contrast specified above, a one-sample t-test was used to identify significant activation on a group level, thresholded using an FDR correction. An F-test thresholded using a Bonferroni correction was also applied. Results are reported in MNI-space, and anatomical labels were derived using the Harvard-Oxford atlas (Notter et al., 2019).

We applied the minimum conjunction procedure to all virtue and vice F-contrasts to identify the global, common loci of activity (Nichols et al., 2005). The resulting regions we call *the core moral network*. We also identified brain regions uniquely involved in specific foundations using T-contrasts that contrasted one vs. the mean of the other 6 moral foundation regressors at the group level.

#### 5.5.2. Inter-subject pattern correlation (ISPC)

To assess how the core moral network represents moral information of the audio narratives, we ran an inter-subject pattern correlation analysis (see Figure 2) on the top clusters of the core moral network (i.e., left and right MTG/STG and right TPJ). This yielded a representation dissimilarity matrix (RDM) of absolute pairwise differences in sentence-level moral–nonmoral ratio scores as provided by the extended MAC dictionary (see Identifying moral foundation values in the stimuli), excluding 0s.

**Figure 2.**
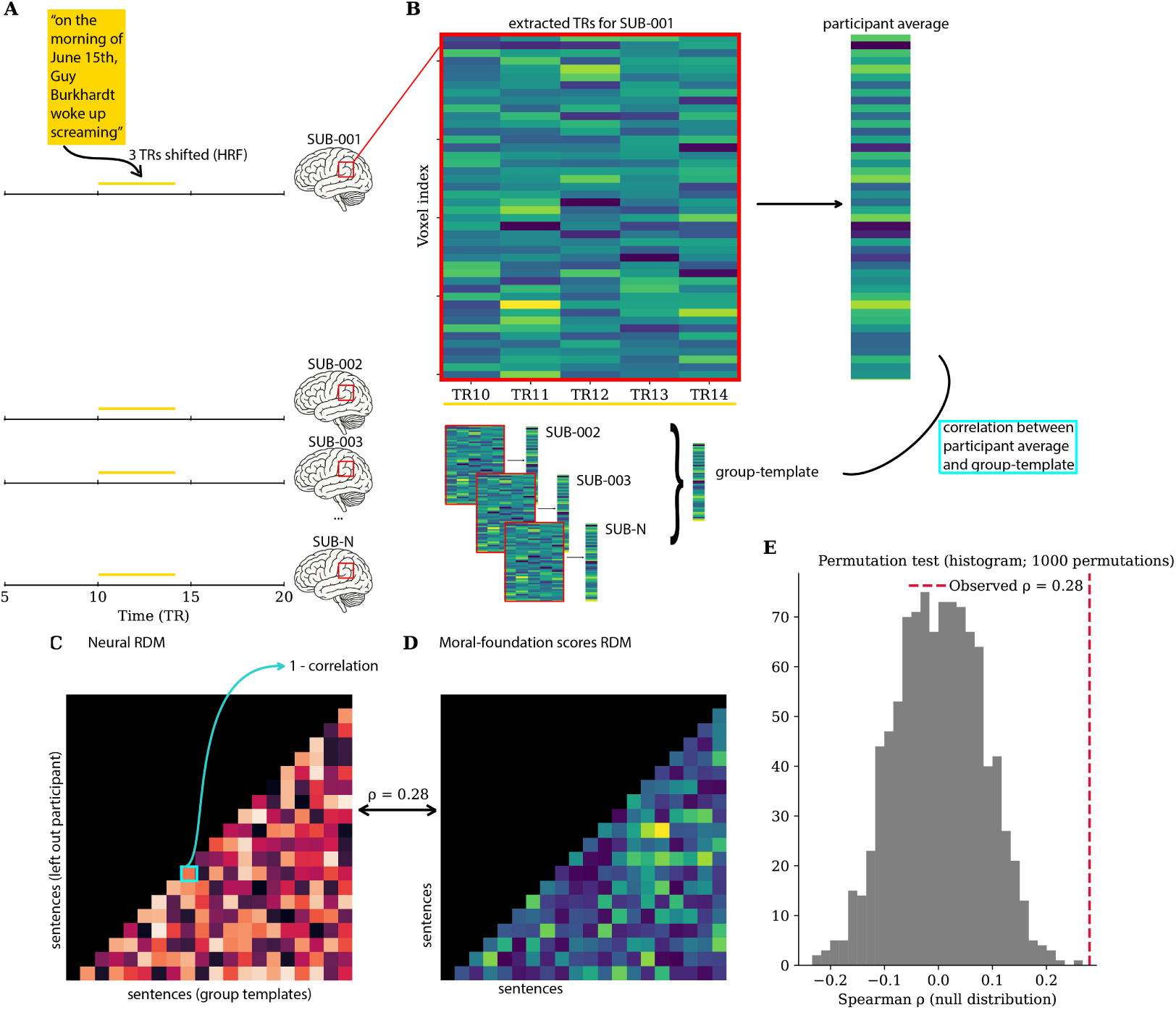
Intersubject pattern correlation analysis logic for comparing the moral-foundation scores RDM to the neural RDMs of the ROIs. [A]. Labeling text to TRs. **[B]** Extracting the relevant voxels for TRs that map onto the sentence to be analyzed, and calculating a participant average neural pattern to be compared to the group average template. **[C]** Neural RDM. **[D]** Text-based RDM. **[E]** Permutation testing to determine significance against the null distribution.

##### Extraction of event-locked multi-voxel activity patterns

For each participant and ROI, we extracted mean multi-voxel activity patterns for each sentence duration. Corresponding event onset and offset TRs were shifted by 3 volumes to account for the HRF.

##### Leave-one-participant-out group templates

inter-subject pattern correlations used a *leave-one-participant-out* method. For each event and each participant, we averaged the patterns of all other participants, getting a constructed participant-specific group-average.

##### Inter-Subject Pattern Correlation Representational Dissimilarity Matrices (ISPC-RDMs)

For each ROI (See ROI selection), we created an ISPC-based RDM. The matrix consisted of Pearson correlations between the individual participant’s neural pattern for sentence i and that participant’s group-average neural pattern for sentence j, for all sentence pairs (i, j). Dissimilarity values were defined as 1 minus the mean correlation across participants. This resulted in a sentence-by-sentence RDM for each ROI.

All RDMs were then vectorized by extracting the upper-triangle values (excluding the diagonal). Correlation between the neural RDM and the moral-content RDM was quantified using Spearman rank correlation, computed separately for each ROI. Significance values were obtained from the associated two-tailed Spearman tests. Significance was determined based on permutation testing (using 1000 permutations), by randomizing the moralcontent RDM and correlating that to the actual neural RDM to generate a null distribution.

### 5.6. ROI selection

We created an anatomically defined TPJ mask using the Harvard-Oxford atlas (Desikan et al., 2006) by taking into account the angular gyrus, supramarginal gyrus, and LOC superior. Subsequently, we intersected this anatomical mask with a meta-analytic mask from neurosynth.org (Yarkoni et al., 2011). The meta-analytic mask was obtained by downloading the association test map for the term “TPJ”, which was thresholded using false discovery rate correction at *q <* 0.01. The map was provided in MNI152 space with 2-mm isotropic resolution. The map was additionally thresholded at *z ≥* 3.0 and binarized. The MTG/STG mask was created using the same procedure. For it, we took the temporal pole, MTG (anterior, posterior, and temporooccipital), STG (anterior, posterior, and temporo-occipital) into account.

## Author CRediT statement

**MK:** Conceptualization, Software, Formal analysis, Investigation, Data Curation, Writing - original draft, Visualization. **SHPC:** Conceptualization, Software, Formal analysis, Investigation, Data Curation, Writing - original draft, Visualization.

## Appendix A. MAC

**Figure A.1.**
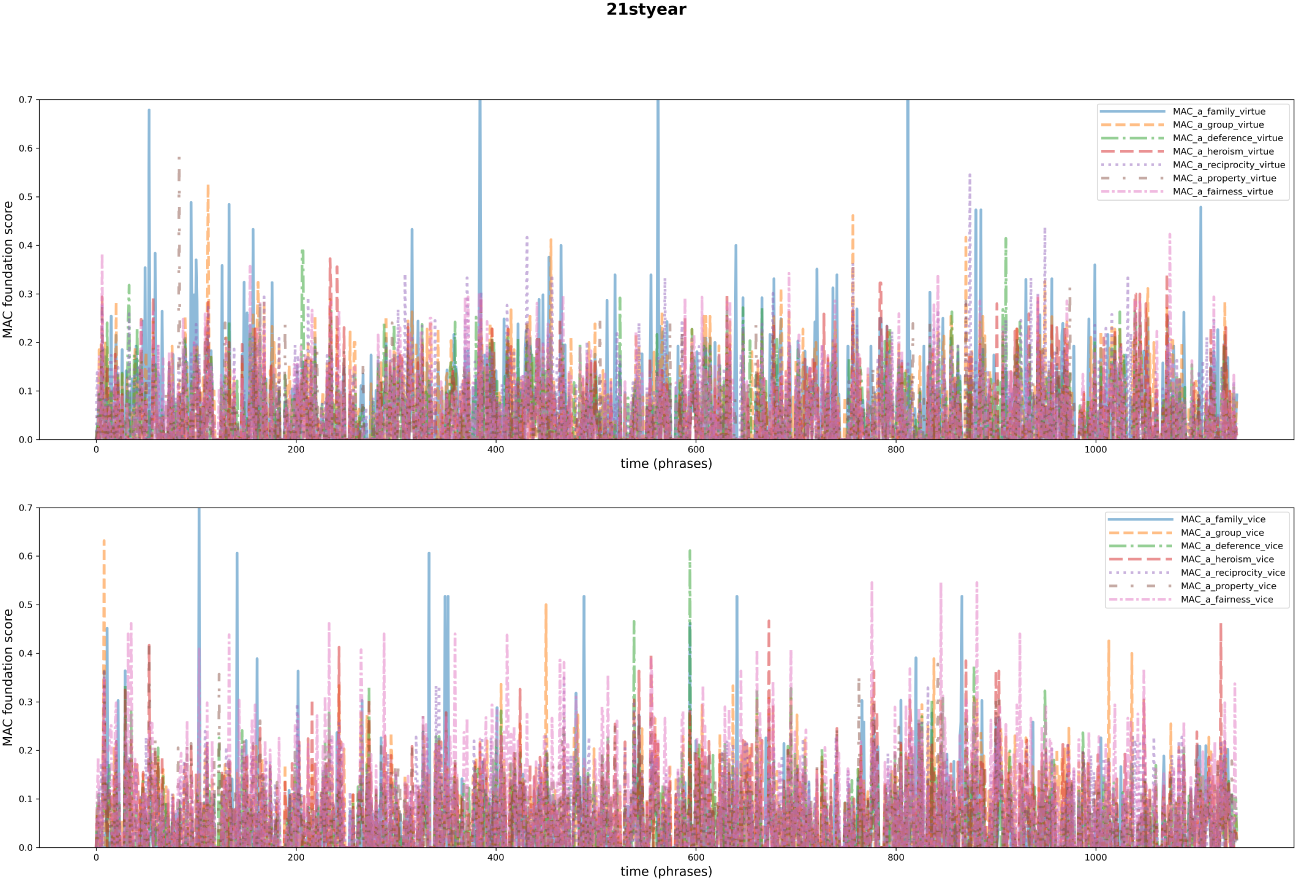
[Top] is 7 MAC virtue domain scores sentence-by-sentence in the 21st Year story. **[Bottom]** is 7 MAC vice domain scores sentence-by-sentence in the 21st Year story.

**Table A.1.** Morality-as-cooperation foundations adopted from Curry et al. (2019).

| Label | Problem | Solution | Virtues | Vices |
| --- | --- | --- | --- | --- |
| Family | Kin selection | Kin Altruism | Duty of care, special obligations to kin | Incest, neglect |
| Group | Coordination | Mutualism | Loyalty, unity, solidarity, conformity | Betrayal, treason |
| Reciprocity | Social Dilemma | Reciprocal Altruism | Reciprocity, trustworthiness, forgiveness | Cheating, ingratitude |
| Heroism | Conflict | Hawkish Displays | Bravery, fortitude, largesse | Cowardice, miserliness |
| Deference | Conflict | Dove-ish Displays | Respect, obedience, humility | Disrespect, hubris |
| Fairness | Conflict | Division | Fairness, impartiality, equality | Unfairness, favouritism |
| Property | Conflict | Ownership | Respect for property, property rights | Theft, trespass |

**Figure A.3.**
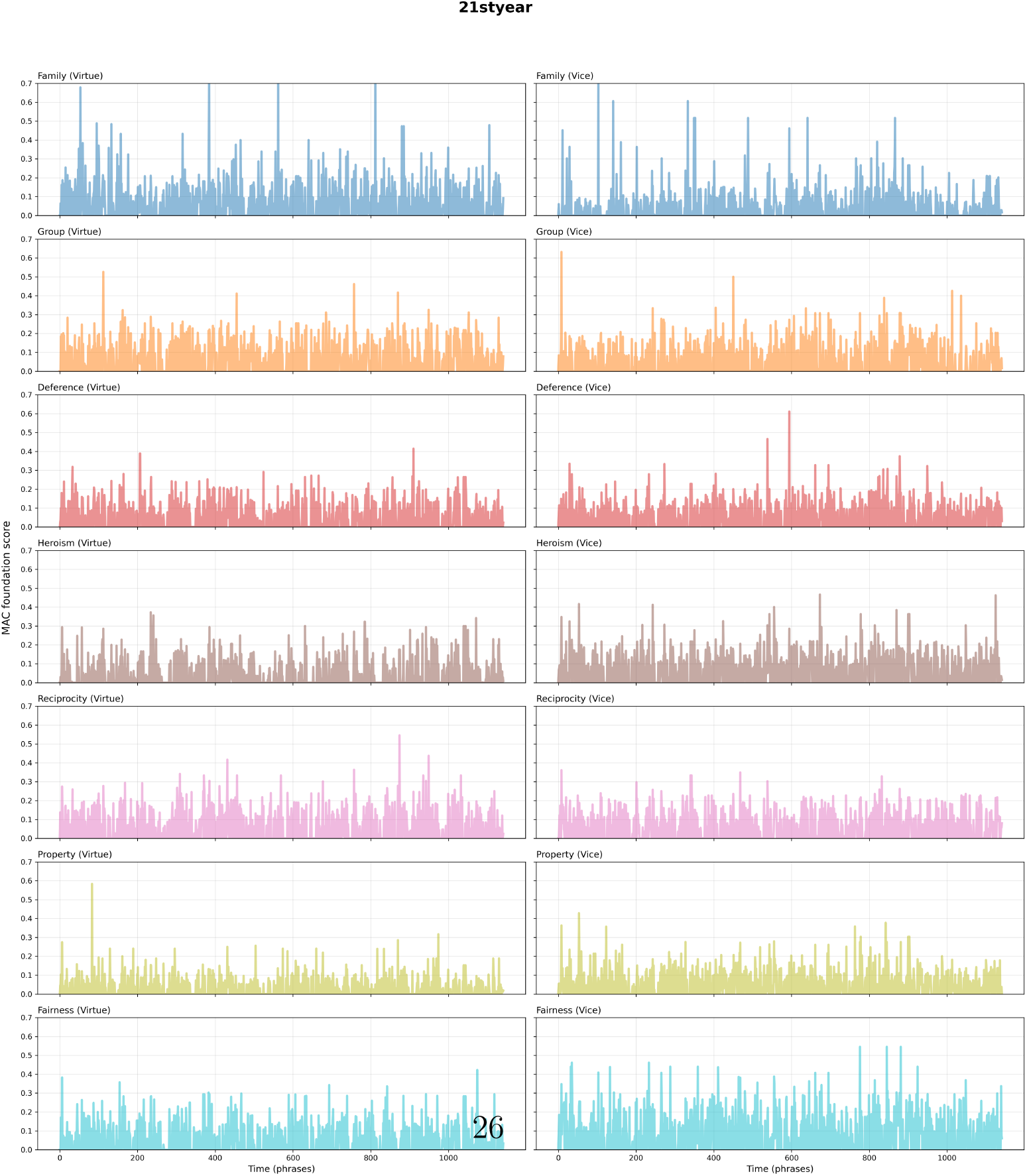
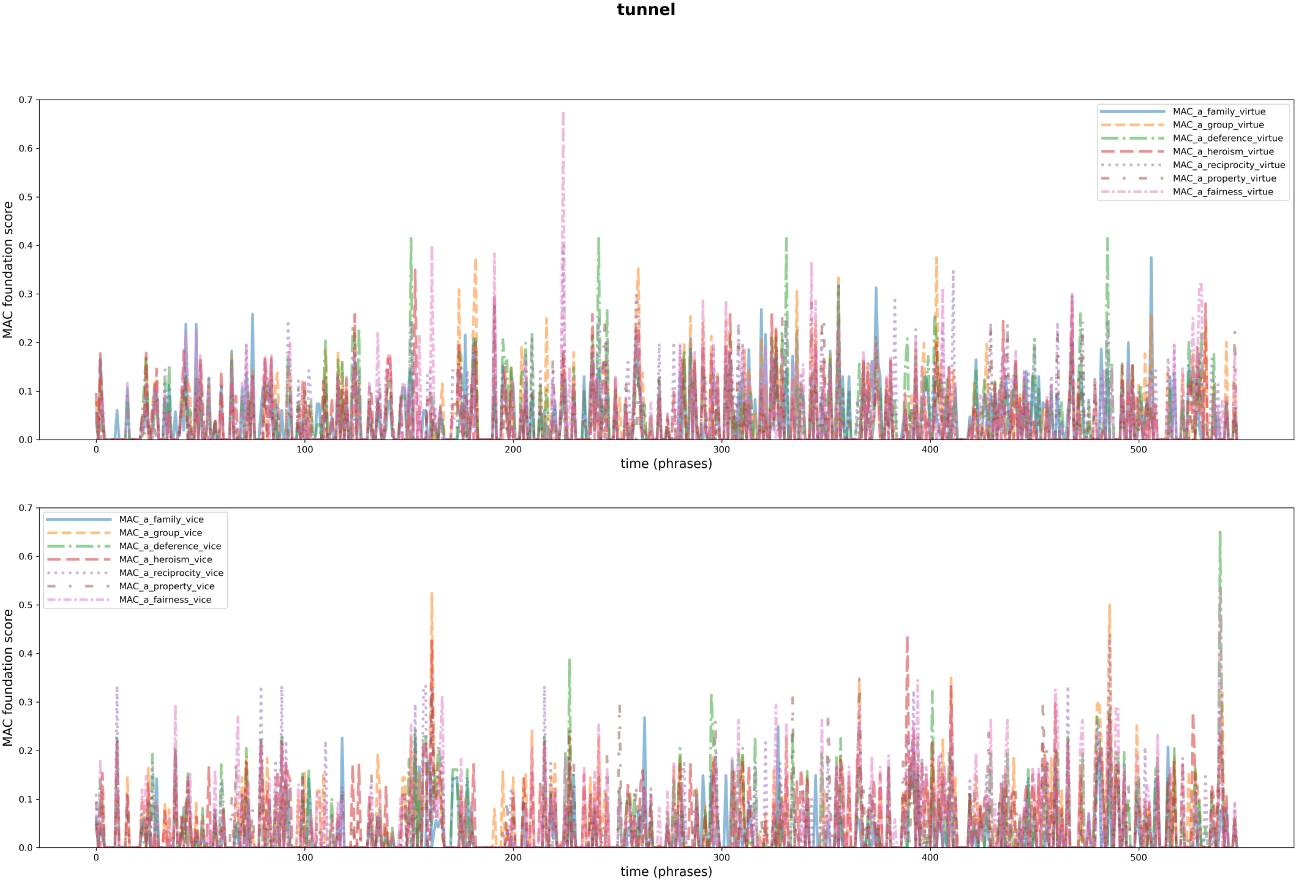
[Top] is 7 MAC virtue domain scores sentence-by-sentence in the Tunnel story. **[Bottom]** is 7 MAC vice domain scores sentence-by-sentence in the Tunnel story.

**Figure A.4.**
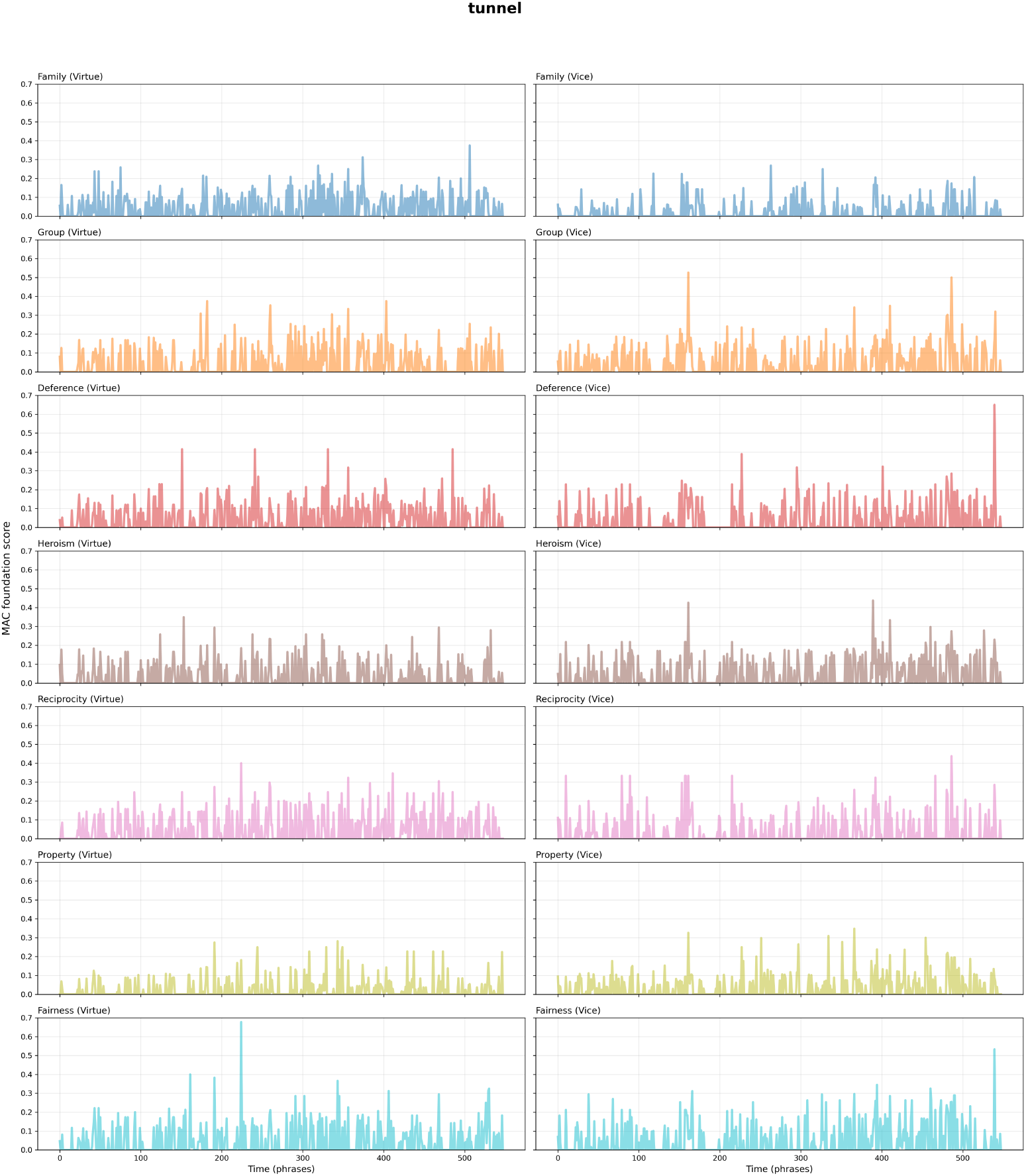
[Left] is 7 MAC virtue domain scores sentence-by-sentence in the Tunnel story, from top to bottom: family, group, deference, heroism, reciprocity, property, fairness. **[Right]** is 7 MAC vice domain scores sentence-by-sentence in the Tunnel story, from top to bottom: family, group, deference, heroism, reciprocity, property, fairness.

## Appendix B. Sentence distribution across the moral domains

**Table B.1.** Distribution of sentences of the Tunnel story across the 7 domains, separately for virtue and vice moral domains.

| Code | Foundation | Virtue (# sentences) | Vice (# sentences) |
| --- | --- | --- | --- |
| 0 | baseline | 248 | 262 |
| 1 | fairness | 61 | 67 |
| 2 | family | 51 | 13 |
| 3 | heroism | 30 | 41 |
| 4 | reciprocity | 62 | 36 |
| 5 | group | 44 | 60 |
| 6 | property | 11 | 30 |
| 7 | deference | 41 | 39 |

**Table B.2.** Distribution of sentences of the 21st Year story across the 7 domains, separately for virtue and vice moral domains.

| Code | Foundation | Virtue (# sentences) | Vice (# sentences) |
| --- | --- | --- | --- |
| 0 | baseline | 366 | 367 |
| 1 | fairness | 112 | 243 |
| 2 | family | 164 | 67 |
| 3 | heroism | 72 | 121 |
| 4 | reciprocity | 119 | 95 |
| 5 | group | 190 | 97 |
| 6 | property | 31 | 65 |
| 7 | deference | 88 | 87 |

**Example sentences: Fairness (virtue):** Did you have to buy a $400 bottle of scotch (0.3830)? **Fairness (vice):** Can’t afford it for one (0.4091). **Family (virtue):** So in love with this baby, spitting up goo on her lap (0.4884). **Family (vice):** He holds a cracker piled with overripe brie and jelly, shoves it in his wife’s mouth (0.6061). **Heroism (virtue):** She wants to protect herself (0.3727). **Heroism (vice):** No. I’ve been afraid to go in (0.4375). **Reciprocity (virtue):** This is crazy, but it fits the facts when I think about it (0.3466). **Reciprocity (vice):** What’s wrong with being upset (0.3333)? **Group (virtue):** You must be so proud of him, Clara (0.5263). **Group (vice):** He seems distracted, guilty of something (0.6316). **Property (virtue):** That your stocks have all tanked (0.5833)? **Property (vice):** He slides over a dirty martini and shrugs off her money (0.3779). **Deference (virtue):** We should have talked this move through before you applied, he laughs (0.3889). **Deference (vice):** Are we broke (0.6111)?

## Appendix C. ISPC results

**Figure C.1.**
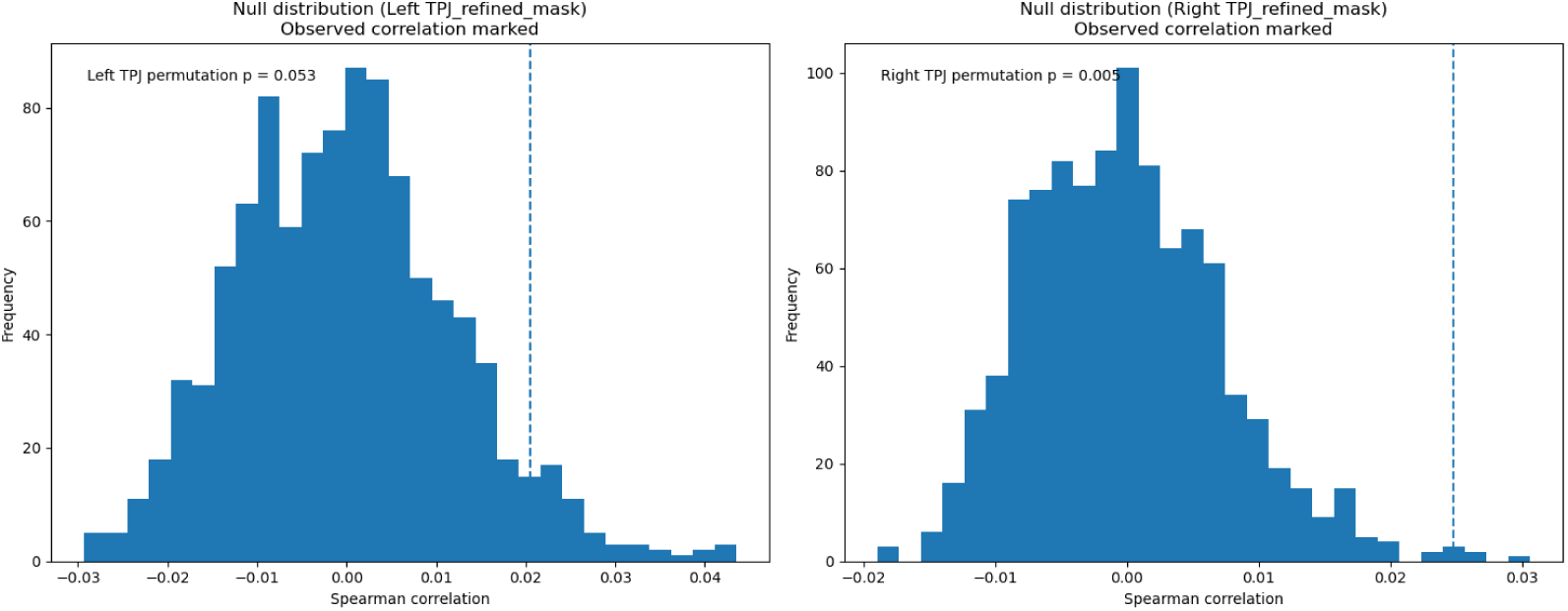
Intersubject pattern correlation (ISPC) results for the TPJ in the Tunnel story. The histogram shows the distribution of Spearman correlation coefficients obtained from 1000 random permutations of the moral-foundation-scores-RDM while correlating with the actual RDM of the left and the right TPJ neural patterns (for the Tunnel story). The vertical dashed line marks the observed correlation in the actual data, indicating that the representational structure in the right TPJ reliably reflects moral similarity between events.

**Figure C.2.**
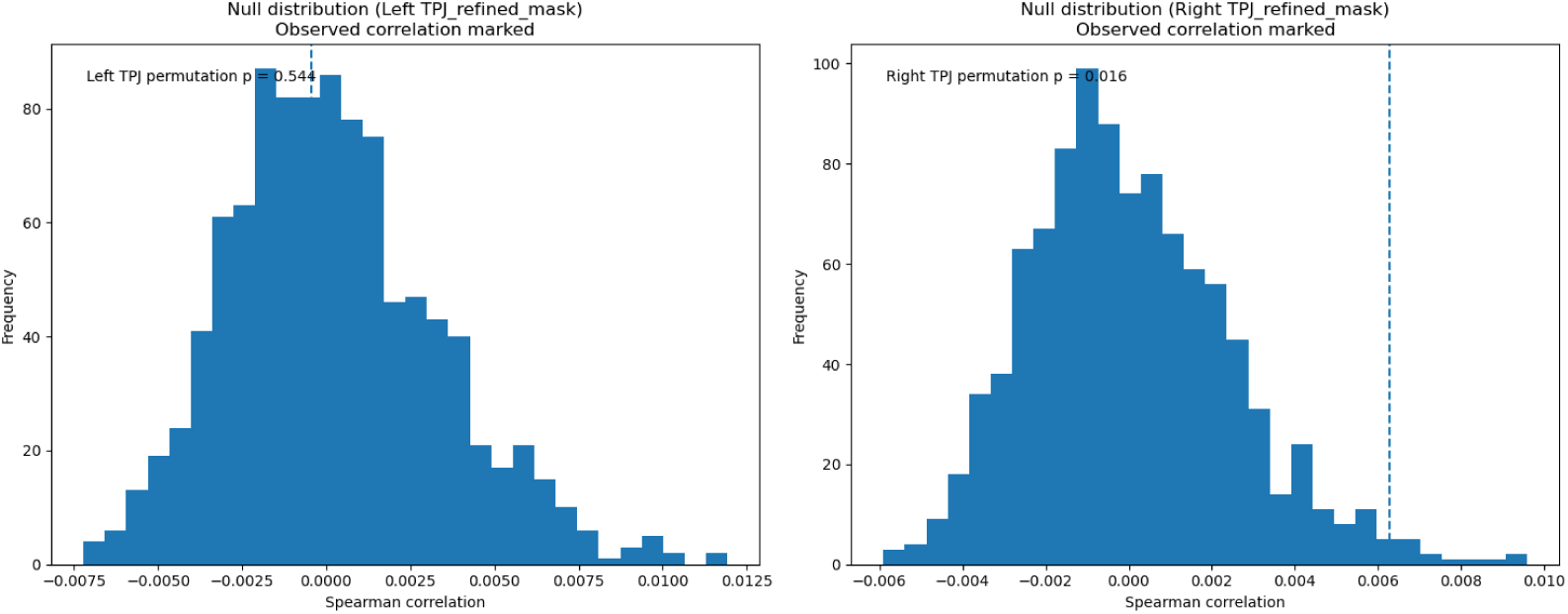
ISPC results for the TPJ in the 21styear story. The same histograms as in figure C.1 with null distribution and a vertical dashed line marking the observed actual correlation for left and right TPJ in the 21st Year story.

**Figure C.3.**
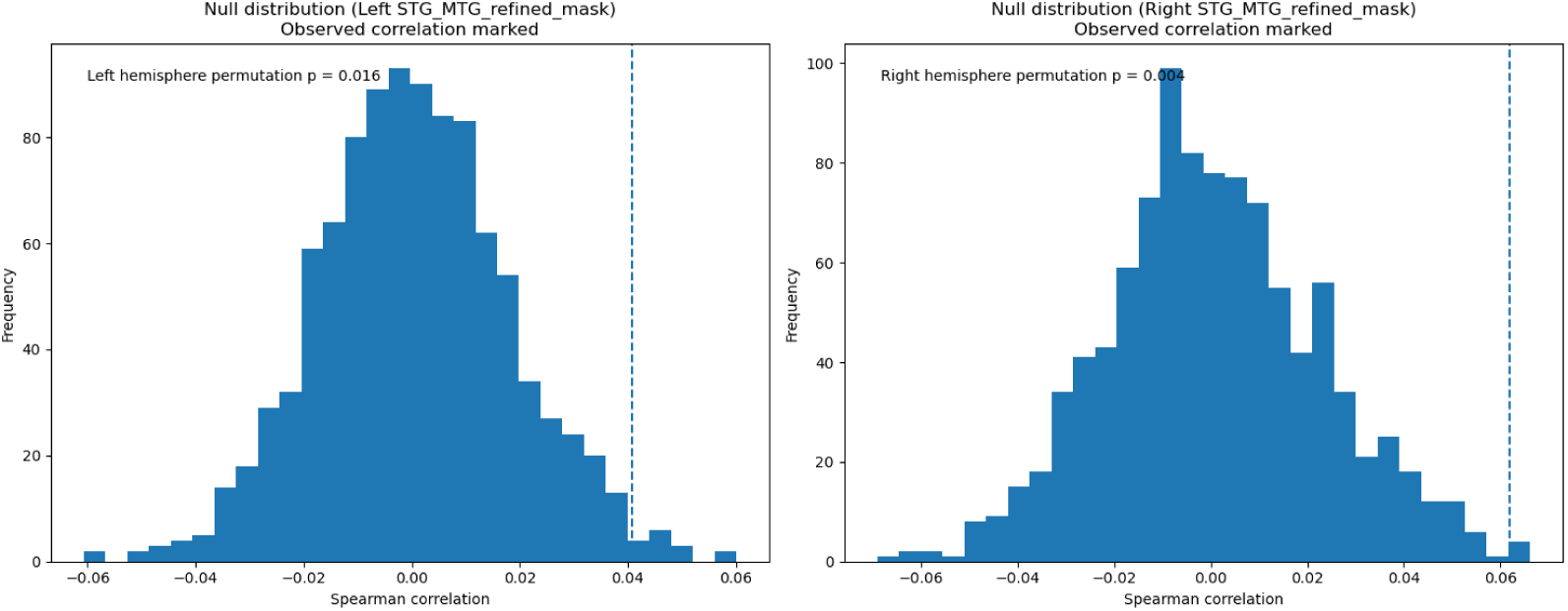
ISPC results for the MTG/STG in the Tunnel story. The histogram shows the distribution of Spearman correlation coefficients obtained from 1000 random permutations of the moral-foundation-scores-RDM while correlating with the actual RDM of the left and the right MTG/STG neural patterns (for the Tunnel story). The vertical dashed line marks the observed correlation in the actual data.

**Figure C.4.**
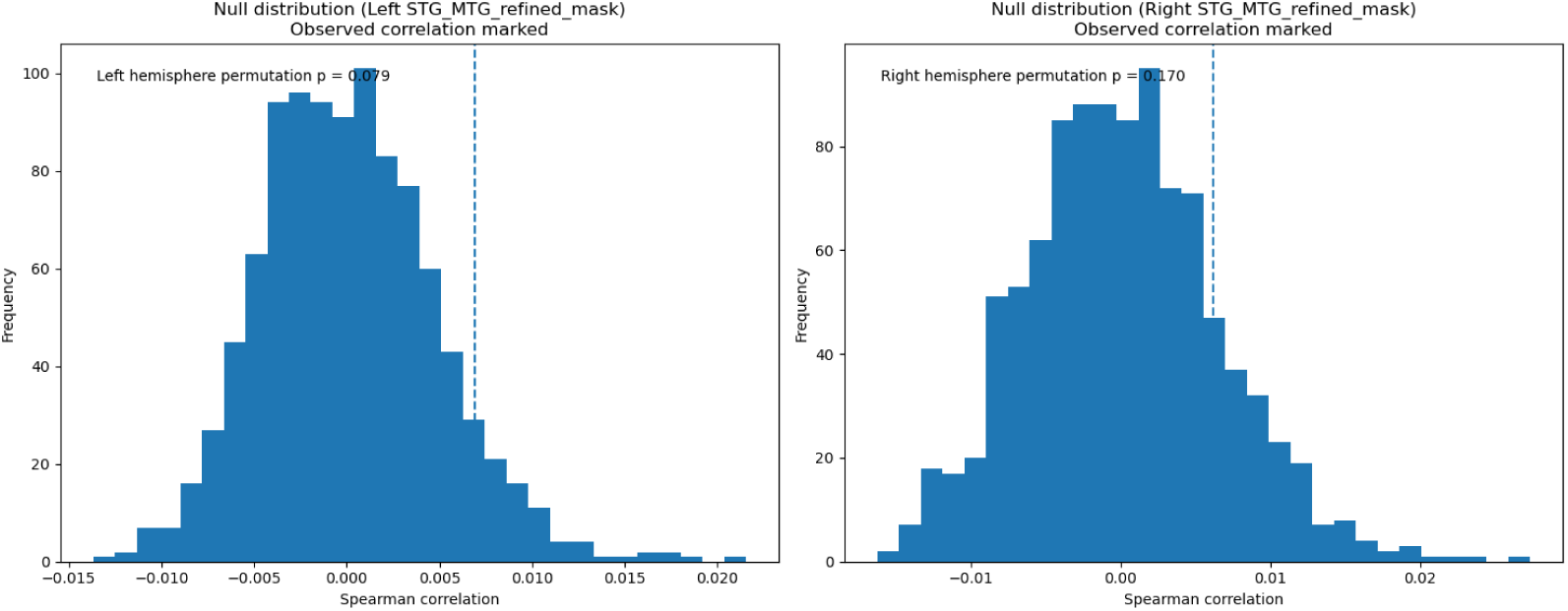
ISPC results for the MTG/STG in the 21styear story. The same histograms as in figure C.3 with null distribution and a vertical dashed line marking the observed actual correlation for left and right MTG/STG in the 21st Year story.

## Appendix D. Details on the foundation-specific brain networks

**Table D.1.** Clusters and atlas labels for fairness virtue for the Tunnel story. Abbreviations: L = left, R = right, INS = insula, FO = frontal operculum, FP = frontal pole, ACC = anterior cingulate cortex, PCG = paracingulate gyrus.

| x | y | z | mean | mm<sup>3</sup> | harvard-oxford labels |
| --- | --- | --- | --- | --- | --- |
| -39.5 | 8.5 | 1.5 | 4.3871 | 324 | 55.56% L INS; 44.44% L FO |
| 35.5 | 50.5 | 29.5 | 4.2573 | 144 | 100.00% R FP |
| -36.5 | 56.5 | 21.5 | 4.3398 | 144 | 100.00% L FP |
| 2.5 | 32.5 | 25.5 | 4.1094 | 108 | 66.67% R ACC; 33.33% R PCG |

**Table D.2.**
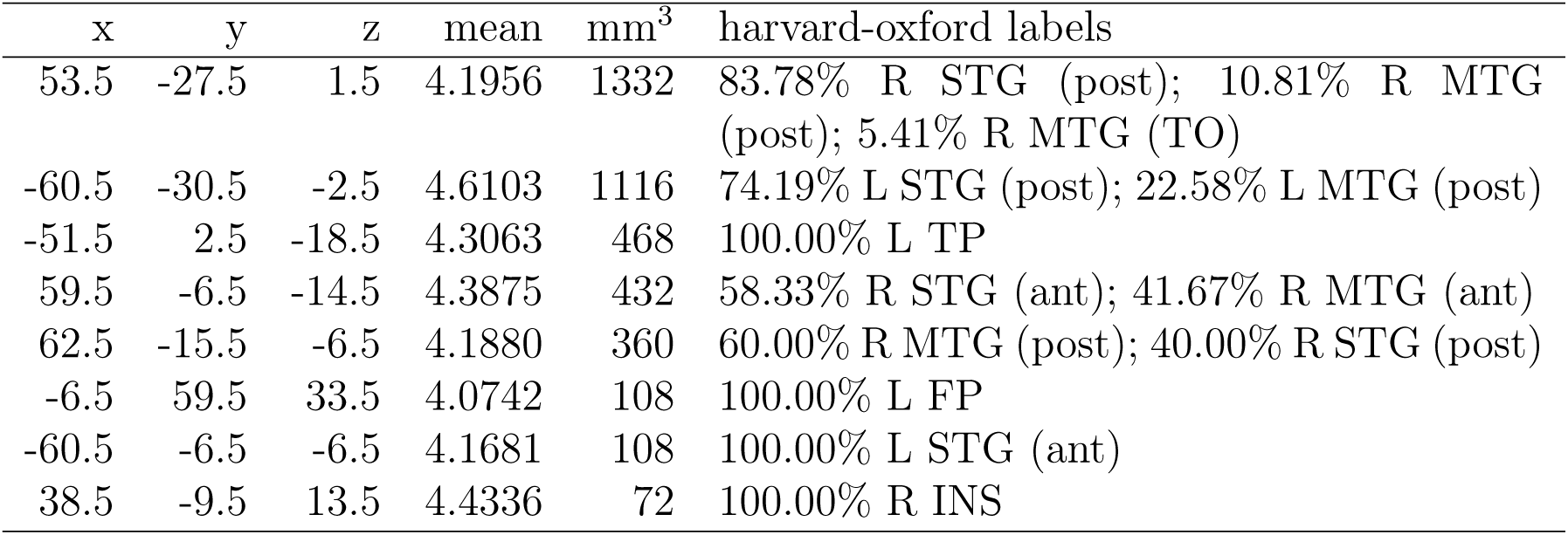
Clusters and atlas labels for family virtue for the 21st Year story. Abbreviations: L = left, R = right, STG = superior temporal gyrus, MTG = middle temporal gyrus, TP = temporal pole, FP = frontal pole, INS = insula, ant = anterior division, post = posterior division, TO = temporo-occipital part.

| x | y | z | mean | mm<sup>3</sup> | harvard-oxford labels |
| --- | --- | --- | --- | --- | --- |
| 53.5 | -27.5 | 1.5 | 4.1956 | 1332 | 83.78% R STG (post); 10.81% R MTG (post); 5.41% R MTG (TO) |
| -60.5 | -30.5 | -2.5 | 4.6103 | 1116 | 74.19% L STG (post); 22.58% L MTG (post) |
| -51.5 | 2.5 | -18.5 | 4.3063 | 468 | 100.00% L TP |
| 59.5 | -6.5 | -14.5 | 4.3875 | 432 | 58.33% R STG (ant); 41.67% R MTG (ant) |
| 62.5 | -15.5 | -6.5 | 4.1880 | 360 | 60.00% R MTG (post); 40.00% R STG (post) |
| -6.5 | 59.5 | 33.5 | 4.0742 | 108 | 100.00% L FP |
| -60.5 | -6.5 | -6.5 | 4.1681 | 108 | 100.00% L STG (ant) |
| 38.5 | -9.5 | 13.5 | 4.4336 | 72 | 100.00% R INS |

**Table D.3.** Clusters and atlas labels for family virtue for the Tunnel story. Abbreviations: L = left, R = right, STG = superior temporal gyrus, MTG = middle temporal gyrus, ITG = inferior temporal gyrus, TP = temporal pole, AG = angular gyrus, SMG = supramarginal gyrus, SFG = superior frontal gyrus, MFG = middle frontal gyrus, IFG (tri) = inferior frontal gyrus pars triangularis, FP = frontal pole, LOC (sup) = lateral occipital cortex superior division, Precuneus = precuneus cortex, PCC = posterior cingulate cortex, Thal = thalamus, PHG = parahippocampal gyrus, PT = planum temporale, ant = anterior division, post = posterior division, TO = temporo-occipital part.

| x | y | z | mean | mm<sup>3</sup> | harvard-oxford labels |
| --- | --- | --- | --- | --- | --- |
| -48.5 | -30.5 | -6.5 | 3.9038 | 9144 | 47.64% L MTG (post); 27.95% L STG (post); 9.06% L MTG (TO); 6.30% L MTG (ant); 6.30% L STG (ant) |
| 50.5 | -15.5 | -6.5 | 3.8154 | 6984 | 39.69% R MTG (post); 17.01% R STG (post); 13.92% R STG (ant); 10.31% R MTG (ant); 6.19% R MTG (TO); 5.67% R TP |
| -42.5 | -54.5 | 29.5 | 3.5436 | 3672 | 56.86% L AG; 33.33% L LOC (sup); 9.80% L SMG (post) |
| 26.5 | 29.5 | 49.5 | 3.7841 | 3132 | 65.52% R SFG; 34.48% R MFG |
| -57.5 | 2.5 | -18.5 | 3.8120 | 1872 | 84.62% L TP; 7.69% L MTG (ant); 7.69% L STG (ant) |
| -12.5 | 59.5 | 21.5 | 3.6015 | 1332 | 81.08% L FP; 18.92% L SFG |
| 53.5 | -51.5 | 21.5 | 3.4563 | 864 | 95.83% R AG |
| -45.5 | -78.5 | 29.5 | 3.6700 | 612 | 100.00% L LOC (sup) |
| 53.5 | 29.5 | 5.5 | 3.6194 | 576 | 100.00% R IFG (tri) |
| -12.5 | -45.5 | 41.5 | 3.6542 | 468 | 84.62% L Precuneus; 15.38% L PCC |
| -3.5 | -51.5 | -38.5 | 3.6362 | 360 | 90.00% no label; 10.00% Brainstem |
| 44.5 | 17.5 | -22.5 | 3.6862 | 288 | 100.00% R TP |
| -30.5 | 26.5 | 45.5 | 3.7287 | 288 | 100.00% L MFG |
| 11.5 | 68.5 | 13.5 | 3.8060 | 288 | 100.00% R FP |
| 5.5 | -60.5 | 45.5 | 3.4363 | 252 | 100.00% R Precuneus |
| -18.5 | 32.5 | 37.5 | 3.5855 | 216 | 100.00% L SFG |
| -18.5 | 17.5 | 45.5 | 3.6058 | 180 | 100.00% L SFG |
| 8.5 | -12.5 | 5.5 | 3.5504 | 180 | 100.00% R Thal |
| 14.5 | -48.5 | 41.5 | 3.2788 | 180 | 100.00% R Precuneus |
| 8.5 | -48.5 | -42.5 | 3.5427 | 144 | 100.00% Brainstem |
| -60.5 | -54.5 | 33.5 | 3.3091 | 144 | 50.00% L AG; 50.00% L SMG (post) |
| 53.5 | -33.5 | 9.5 | 3.5181 | 144 | 50.00% R SMG (post); 50.00% R STG (post) |
| -21.5 | 14.5 | 53.5 | 3.4364 | 108 | 100.00% L SFG |
| 44.5 | -45.5 | 29.5 | 3.6287 | 108 | 100.00% R AG |
| -9.5 | 11.5 | 65.5 | 3.6227 | 72 | 100.00% L SFG |
| -9.5 | -30.5 | 5.5 | 4.4109 | 72 | 100.00% L Thal |
| 56.5 | -24.5 | 5.5 | 3.4778 | 72 | 50.00% R PT; 50.00% R STG (post) |
| 20.5 | -27.5 | -18.5 | 3.5285 | 72 | 100.00% R PHG (post) |
| 38.5 | -78.5 | 41.5 | 3.3170 | 72 | 100.00% R LOC (sup) |
| 41.5 | 5.5 | 49.5 | 3.3636 | 72 | 100.00% R MFG |
| 47.5 | 2.5 | -46.5 | 3.3845 | 72 | 100.00% R ITG (ant) |
| 59.5 | -54.5 | 17.5 | 3.3644 | 72 | 100.00% R AG |

**Table D.4.** Clusters and atlas labels for group virtue for the Tunnel story. Abbreviations: R = right, MTG = middle temporal gyrus, TO = temporo-occipital part.

| x | y | z | mean | mm<sup>3</sup> | harvard-oxford labels |
| --- | --- | --- | --- | --- | --- |
| 35.5 | -54.5 | 9.5 | 4.0159 | 72 | 50.00% R MTG (TO); 50.00% no label |

**Table D.5.** Clusters and atlas labels for group virtue for the 21st Year story. Abbreviations: L = left, R = right, IFG (op) = inferior frontal gyrus pars opercularis, SFG = superior frontal gyrus, FP = frontal pole, LatVent = lateral ventricle.

| x | y | z | mean | mm<sup>3</sup> | harvard-oxford labels |
| --- | --- | --- | --- | --- | --- |
| 11.5 | 5.5 | 17.5 | 3.8941 | 648 | 77.78% R Caudate; 22.22% R LatVent |
| -12.5 | 5.5 | 17.5 | 4.3485 | 288 | 75.00% L Caudate; 25.00% L LatVent |
| 53.5 | 17.5 | 5.5 | 3.9747 | 108 | 100.00% R IFG (op) |
| 11.5 | 26.5 | 57.5 | 3.6741 | 72 | 100.00% R SFG |
| -48.5 | 41.5 | 17.5 | 3.8639 | 72 | 100.00% L FP |

**Table D.6.** Clusters and atlas labels for property virtue for the 21st Year story. Abbreviations: L = left, R = right, LOC (sup) = lateral occipital cortex superior division, TOFC = temporal occipital fusiform cortex, TFC (post) = temporal fusiform cortex posterior division, Insula = insular cortex.

| x | y | z | mean | mm<sup>3</sup> | harvard-oxford labels |
| --- | --- | --- | --- | --- | --- |
| -36.5 | -78.5 | 25.5 | 4.0770 | 396 | 100.00% L LOC (sup) |
| -42.5 | -39.5 | 25.5 | 4.2354 | 180 | 100.00% L Parietal Operculum |
| -27.5 | -24.5 | 49.5 | 4.0628 | 144 | 100.00% L Precentral Gyrus |
| 29.5 | 26.5 | 5.5 | 4.0983 | 108 | 100.00% R Insula |
| 26.5 | -39.5 | -18.5 | 3.8447 | 72 | 50.00% R TOFC; 50.00% R TFC (post) |
| 20.5 | -69.5 | 57.5 | 4.0251 | 72 | 100.00% R LOC (sup) |
| 8.5 | -27.5 | 69.5 | 4.0363 | 72 | 100.00% R Precentral Gyrus |
| -9.5 | -48.5 | -2.5 | 3.8885 | 72 | 100.00% L Lingual Gyrus |

**Table D.7.**
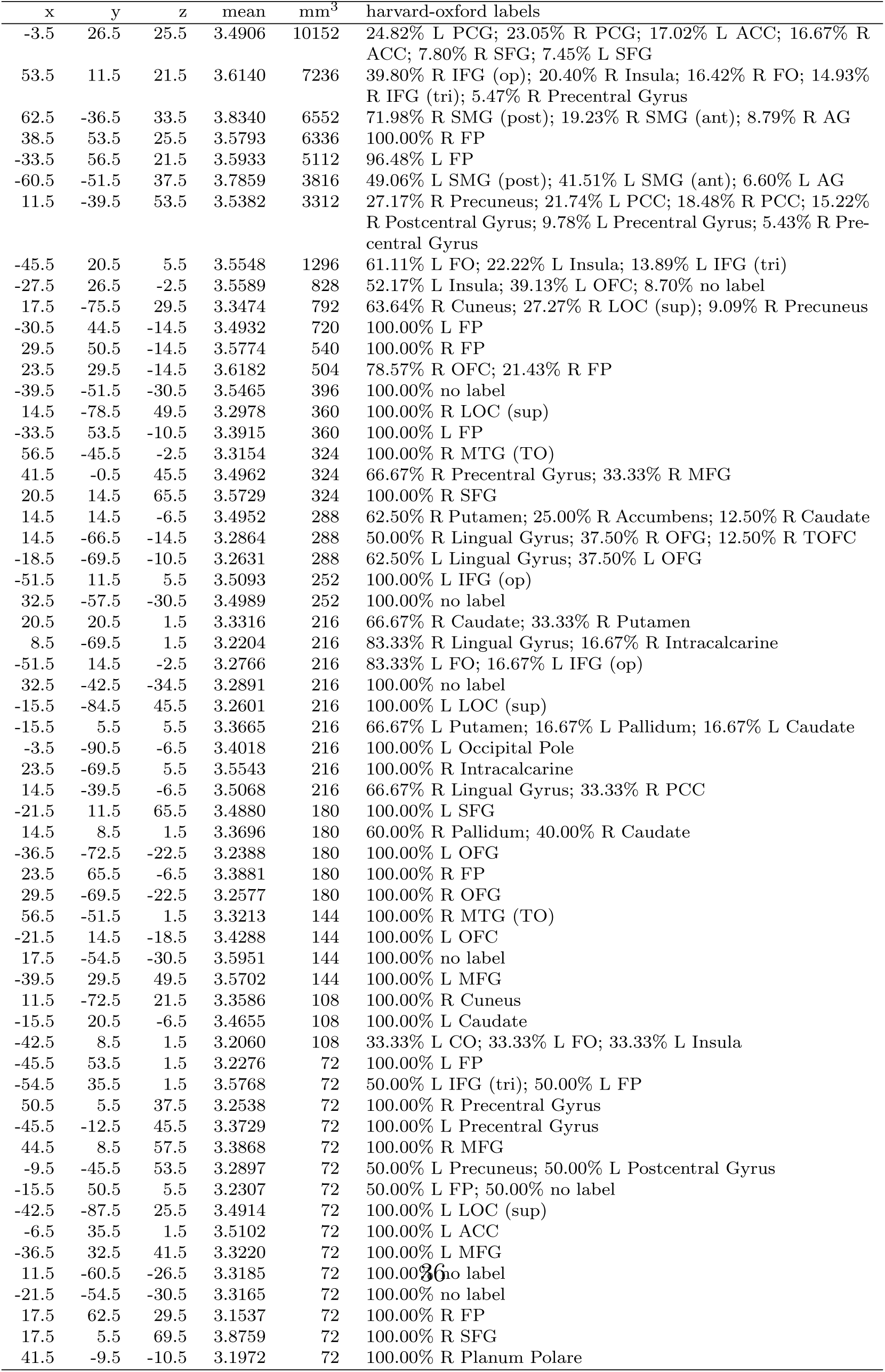
Clusters and atlas labels for reciprocity virtue for the 21st Year story. Abbreviations: L = left, R = right, ACC = anterior cingulate cortex, PCC = posterior cingulate cortex, PCG = paracingulate gyrus, SFG = superior frontal gyrus, MFG =

**Table D.8.** Clusters and atlas labels for fairness vice for the Tunnel story. Abbreviations: L = left, R = right, STG = superior temporal gyrus, SMG = supramarginal gyrus, PT = planum temporale, HG = Heschl’s gyrus, TP = temporal pole, PHG = parahippocampal gyrus, Thal = thalamus, CO = central operculum, ant = anterior division, post = posterior division.

| x | y | z | mean | mm<sup>3</sup> | harvard-oxford labels |
| --- | --- | --- | --- | --- | --- |
| 50.5 | -18.5 | 9.5 | 4.0766 | 7560 | 29.52% R STG (post); 27.14% R PT; 17.62% R HG; 16.19% R SMG (post) |
| -63.5 | -30.5 | 13.5 | 4.1146 | 612 | 58.82% L STG (post); 41.18% L PT |
| 20.5 | -6.5 | -14.5 | 4.2835 | 612 | 100.00% R Amygdala |
| -48.5 | -0.5 | -6.5 | 4.1650 | 468 | 69.23% L Planum Polare; 23.08% L TP; 7.69% L CO |
| -60.5 | -15.5 | 5.5 | 4.0838 | 396 | 100.00% L PT |
| 29.5 | -0.5 | -22.5 | 3.9304 | 360 | 100.00% R Amygdala |
| 50.5 | 17.5 | -22.5 | 4.0404 | 252 | 100.00% R TP |
| -57.5 | -48.5 | 13.5 | 3.8366 | 108 | 100.00% L SMG (post) |
| -48.5 | -21.5 | 5.5 | 3.8456 | 108 | 100.00% L HG |
| 26.5 | -30.5 | 17.5 | 3.9343 | 108 | 100.00% no label |
| 68.5 | -27.5 | 17.5 | 3.7942 | 108 | 66.67% R STG (post); 33.33% R PT |
| 62.5 | 5.5 | -6.5 | 3.7109 | 108 | 66.67% R STG (ant); 33.33% R TP |
| 20.5 | -15.5 | 17.5 | 3.7697 | 72 | 50.00% R Thal; 50.00% no label |
| 23.5 | -75.5 | -30.5 | 3.8267 | 72 | 100.00% no label |
| 26.5 | 5.5 | -30.5 | 3.8127 | 72 | 50.00% R PHG (ant); 50.00% R TP |
| 47.5 | 11.5 | -14.5 | 4.2680 | 72 | 100.00% R TP |
| 68.5 | -42.5 | 13.5 | 3.7079 | 72 | 100.00% R SMG (post) |

**Table D.9.** Clusters and atlas labels for family vice for the Tunnel story. Abbreviations: L = left, R = right, LOC (sup) = lateral occipital cortex superior division, PCC = posterior cingulate cortex, AG = angular gyrus.

| x | y | z | mean | mm<sup>3</sup> | harvard-oxford labels |
| --- | --- | --- | --- | --- | --- |
| 11.5 | -54.5 | 21.5 | 3.8131 | 684 | 100.00% R Precuneus |
| -15.5 | -57.5 | 17.5 | 3.7261 | 324 | 100.00% L Precuneus |
| 41.5 | -72.5 | 33.5 | 3.7931 | 216 | 100.00% R LOC (sup) |
| 17.5 | -45.5 | 37.5 | 3.6856 | 180 | 60.00% R PCC; 40.00% R Precuneus |
| 50.5 | -51.5 | 33.5 | 4.0294 | 72 | 100.00% R AG |

**Table D.10.**
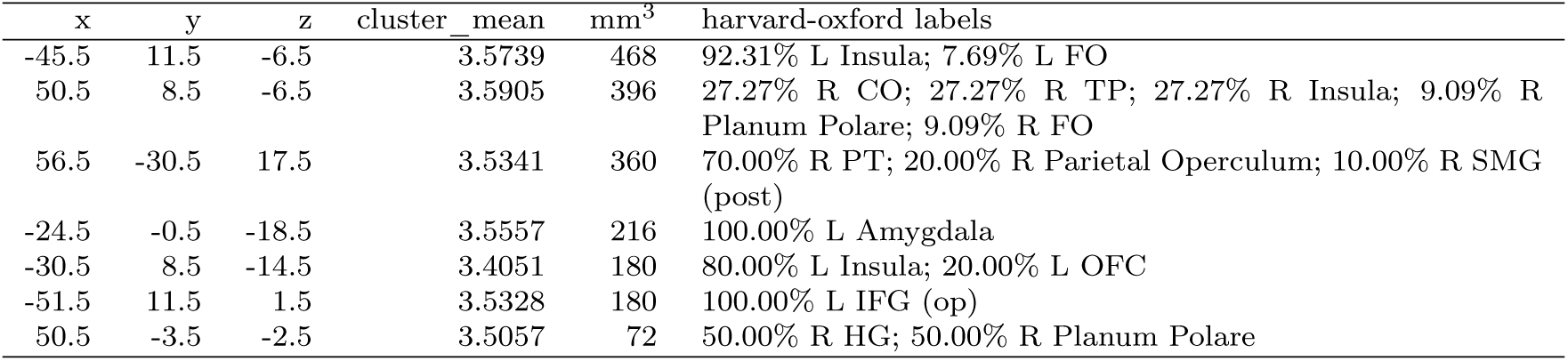
Clusters and atlas labels for group vice for the Tunnel story. Abbreviations: L = left, R = right, Insula = insular cortex, FO = frontal operculum, CO = central operculum, TP = temporal pole, PT = planum temporale, HG = Heschl’s gyrus, SMG = supramarginal gyrus, IFG (op) = inferior frontal gyrus pars opercularis, OFC = orbitofrontal cortex.

**Table D.11.** Clusters and atlas labels for reciprocity vice for the Tunnel story. Abbreviations: L = left, R = right, STG = superior temporal gyrus, MTG = middle temporal gyrus, SMG = supramarginal gyrus, AG = angular gyrus, PT = planum temporale, ant = anterior division, post = posterior division, TO = temporo-occipital part.

| x | y | z | mean | mm<sup>3</sup> | harvard-oxford labels |
| --- | --- | --- | --- | --- | --- |
| 62.5 | -18.5 | -2.5 | 4.3453 | 1620 | 75.56% R STG (post); 17.78% R MTG (post) |
| -63.5 | -18.5 | 1.5 | 4.2603 | 756 | 80.95% L STG (post); 19.05% L PT |
| -39.5 | -36.5 | 13.5 | 4.1102 | 180 | 100.00% L PT |
| 50.5 | -36.5 | 9.5 | 4.2007 | 108 | 100.00% R SMG (post) |
| -54.5 | -33.5 | 9.5 | 4.0909 | 108 | 66.67% L PT; 33.33% L STG (post) |
| -66.5 | -36.5 | 13.5 | 4.1243 | 108 | 100.00% L STG (post) |
| 56.5 | -0.5 | -14.5 | 4.1278 | 72 | 100.00% R STG (ant) |
| -57.5 | -42.5 | 1.5 | 4.0187 | 72 | 50.00% L MTG (TO); 50.00% L MTG (post) |
| -63.5 | -51.5 | 17.5 | 4.0144 | 72 | 50.00% L AG; 50.00% L SMG (post) |

**Table D.12.** Clusters and atlas labels for fairness vice for the 21st Year story. Abbreviations: L = left, R = right, STG = superior temporal gyrus, FP = frontal pole, ant = anterior division.

| x | y | z | mean | mm<sup>3</sup> | harvard-oxford labels |
| --- | --- | --- | --- | --- | --- |
| -60.5 | -6.5 | -2.5 | 4.5379 | 144 | 100.00% L STG (ant) |
| -0.5 | 44.5 | -18.5 | 4.7709 | 72 | 100.00% R Medial Frontal Cortex |
| -3.5 | 53.5 | -14.5 | 4.2888 | 72 | 50.00% L Medial Frontal Cortex; 50.00% L FP |

**Table D.13.** Cluster peaks and atlas labels for family vice for the 21st Year story. Abbreviations: L = left, R = right, STG = superior temporal gyrus, PT = planum temporale, HG = Heschl’s gyrus, CO = central operculum, ant = anterior division, post = posterior division.

| x | y | z | mean | mm<sup>3</sup> | harvard-oxford labels |
| --- | --- | --- | --- | --- | --- |
| 53.5 | -3.5 | -2.5 | 4.3133 | 684 | 36.84% R PT; 31.58% R HG; 10.53% R Planum Polare; 10.53% R CO; 5.26% R STG (post); 5.26% R STG (ant) |
| -60.5 | -15.5 | 1.5 | 4.3309 | 216 | 100.00% L PT |
| -63.5 | -27.5 | 9.5 | 4.4521 | 216 | 50.00% L PT; 50.00% L STG (post) |
| 59.5 | -30.5 | 9.5 | 4.3165 | 180 | 60.00% R STG (post); 40.00% R PT |
| -39.5 | -33.5 | 13.5 | 4.4535 | 180 | 100.00% L PT |

**Table D.14.**
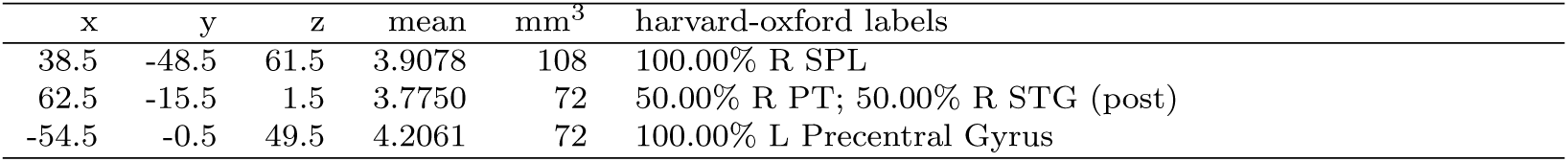
Cluster peaks and atlas labels for group vice for the 21st Year story. Abbreviations: L = left, R = right, SPL = superior parietal lobule, STG = superior temporal gyrus, PT = planum temporale, post = posterior division.

| x | y | z | mean | mm<sup>3</sup> | harvard-oxford labels |
| --- | --- | --- | --- | --- | --- |
| 38.5 | -48.5 | 61.5 | 3.9078 | 108 | 100.00% R SPL |
| 62.5 | -15.5 | 1.5 | 3.7750 | 72 | 50.00% R PT; 50.00% R STG (post) |
| -54.5 | -0.5 | 49.5 | 4.2061 | 72 | 100.00% L Precentral Gyrus |

**Table D.15.** Clusters and atlas labels for heroism vice for the 21st Year story. Abbreviations: L = left, R = right, FP = frontal pole, HG = Heschl’s gyrus, PT = planum temporale, CO = central operculum, MTG = middle temporal gyrus, ant = anterior division, post = posterior division.

| x | y | z | mean | mm<sup>3</sup> | harvard-oxford labels |
| --- | --- | --- | --- | --- | --- |
| 50.5 | -0.5 | 1.5 | 4.2370 | 1764 | 48.98% R CO; 26.53% R HG; 14.29% R Planum Polare;<br>10.20% R PT |
| -3.5 | 59.5 | -10.5 | 4.2047 | 1188 | 66.67% L FP; 18.18% L Medial Frontal Cortex; 9.09% R FP;<br>6.06% R Medial Frontal Cortex |
| -3.5 | 62.5 | 17.5 | 4.0753 | 324 | 100.00% L FP |
| 41.5 | -21.5 | 9.5 | 4.3333 | 144 | 100.00% R HG |
| 35.5 | -33.5 | 17.5 | 4.0066 | 144 | 50.00% R PT; 50.00% R Parietal Operculum |
| -12.5 | 56.5 | 37.5 | 4.1868 | 144 | 100.00% L FP |
| -54.5 | -18.5 | 5.5 | 3.9642 | 144 | 75.00% L HG; 25.00% L CO |
| -63.5 | -12.5 | -18.5 | 3.8682 | 108 | 66.67% L MTG (post); 33.33% L MTG (ant) |

**Table D.16.**
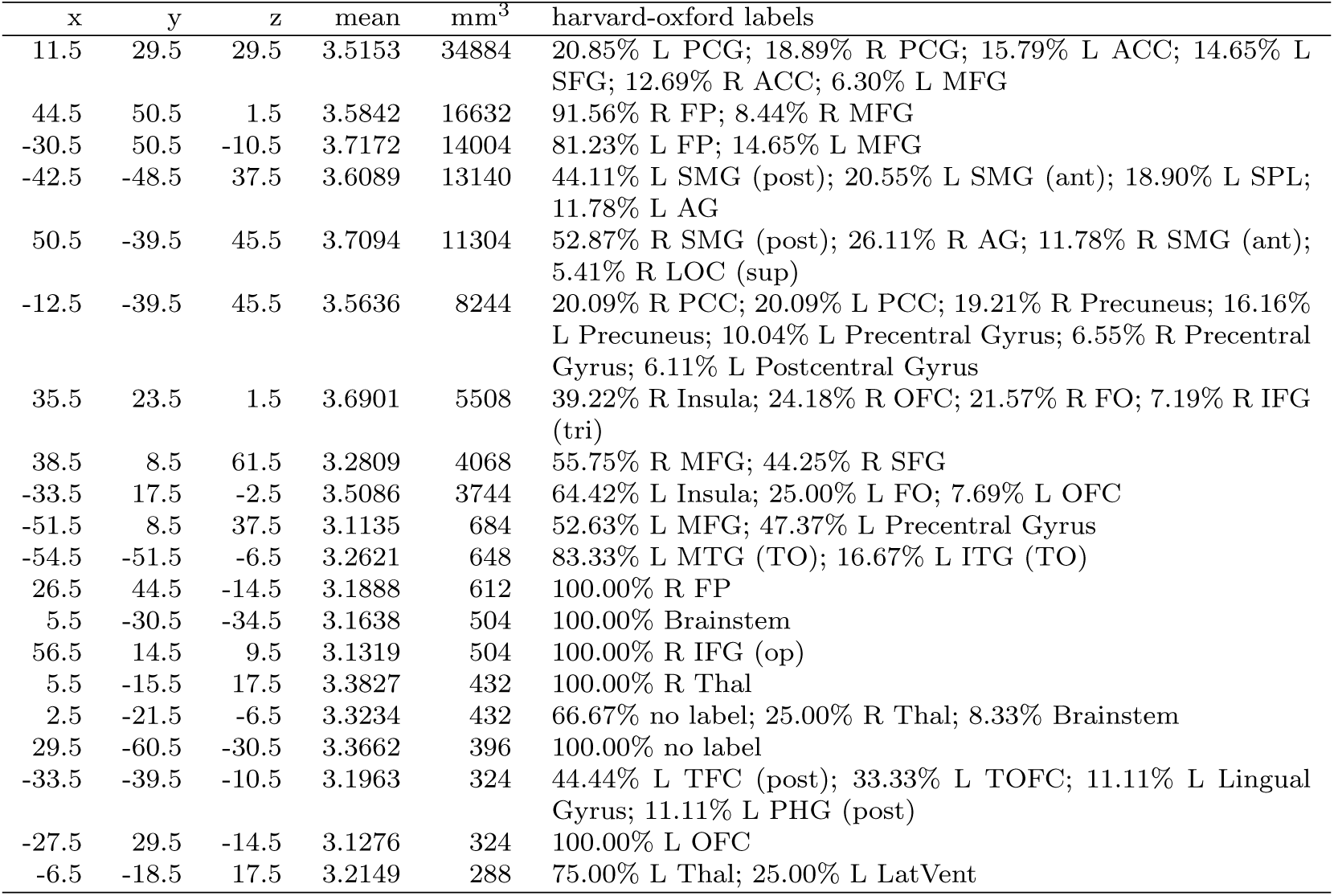
Clusters and atlas labels for property vice for the 21st Year story. Abbreviations: L = left, R = right, PCG = paracingulate gyrus, ACC = anterior cingulate cortex, PCC = posterior cingulate cortex, SFG = superior frontal gyrus, MFG = middle frontal gyrus, FP = frontal pole, SMG = supramarginal gyrus, AG = angular gyrus, SPL = superior parietal lobule, LOC (sup) = lateral occipital cortex superior division, OFC = orbitofrontal cortex, FO = frontal operculum, IFG (op/tri) = inferior frontal gyrus pars opercularis/triangularis, MTG/ITG (TO) = middle/inferior temporal gyrus temporo-occipital part, TFC/TOFC = temporal fusiform cortex / temporal occipital fusiform cortex, PHG = parahippocampal gyrus, Thal = thalamus, LatVent = lateral ventricle.

| x | y | z | mean | mm<sup>3</sup> | harvard-oxford labels |
| --- | --- | --- | --- | --- | --- |
| 11.5 | 29.5 | 29.5 | 3.5153 | 34884 | 20.85% L PCG; 18.89% R PCG; 15.79% L ACC; 14.65% L SFG; 12.69% R ACC; 6.30% L MFG |
| 44.5 | 50.5 | 1.5 | 3.5842 | 16632 | 91.56% R FP; 8.44% R MFG |
| -30.5 | 50.5 | -10.5 | 3.7172 | 14004 | 81.23% L FP; 14.65% L MFG |
| -42.5 | -48.5 | 37.5 | 3.6089 | 13140 | 44.11% L SMG (post); 20.55% L SMG (ant); 18.90% L SPL; 11.78% L AG |
| 50.5 | -39.5 | 45.5 | 3.7094 | 11304 | 52.87% R SMG (post); 26.11% R AG; 11.78% R SMG (ant); 5.41% R LOC (sup) |
| -12.5 | -39.5 | 45.5 | 3.5636 | 8244 | 20.09% R PCC; 20.09% L PCC; 19.21% R Precuneus; 16.16% L Precuneus; 10.04% L Precentral Gyrus; 6.55% R Precentral Gyrus; 6.11% L Postcentral Gyrus |
| 35.5 | 23.5 | 1.5 | 3.6901 | 5508 | 39.22% R Insula; 24.18% R OFC; 21.57% R FO; 7.19% R IFG (tri) |
| 38.5 | 8.5 | 61.5 | 3.2809 | 4068 | 55.75% R MFG; 44.25% R SFG |
| -33.5 | 17.5 | -2.5 | 3.5086 | 3744 | 64.42% L Insula; 25.00% L FO; 7.69% L OFC |
| -51.5 | 8.5 | 37.5 | 3.1135 | 684 | 52.63% L MFG; 47.37% L Precentral Gyrus |
| -54.5 | -51.5 | -6.5 | 3.2621 | 648 | 83.33% L MTG (TO); 16.67% L ITG (TO) |
| 26.5 | 44.5 | -14.5 | 3.1888 | 612 | 100.00% R FP |
| 5.5 | -30.5 | -34.5 | 3.1638 | 504 | 100.00% Brainstem |
| 56.5 | 14.5 | 9.5 | 3.1319 | 504 | 100.00% R IFG (op) |
| 5.5 | -15.5 | 17.5 | 3.3827 | 432 | 100.00% R Thal |
| 2.5 | -21.5 | -6.5 | 3.3234 | 432 | 66.67% no label; 25.00% R Thal; 8.33% Brainstem |
| 29.5 | -60.5 | -30.5 | 3.3662 | 396 | 100.00% no label |
| -33.5 | -39.5 | -10.5 | 3.1963 | 324 | 44.44% L TFC (post); 33.33% L TOFC; 11.11% L Lingual Gyrus; 11.11% L PHG (post) |
| -27.5 | 29.5 | -14.5 | 3.1276 | 324 | 100.00% L OFC |
| -6.5 | -18.5 | 17.5 | 3.2149 | 288 | 75.00% L Thal; 25.00% L LatVent |

## Notes

### Competing Interest Statement

The authors have declared no competing interest.

